# Species richness determines *C. difficile* invasion outcome in synthetic human gut communities

**DOI:** 10.1101/2021.03.23.436677

**Authors:** Susan Hromada, Ryan L. Clark, Yili Qian, Lauren Watson, Nasia Safdar, Ophelia S. Venturelli

## Abstract

Understanding the principles of colonization resistance of the gut microbiome to the pathogen *Clostridioides difficile* will enable the design of next generation defined bacterial therapeutics. We investigate the ecological principles of community resistance to *C. difficile* invasion using a diverse synthetic human gut microbiome. Our results show that species richness is a key determinant of *C. difficile* growth across a wide range of ecological contexts. Using a dynamic computational model, we demonstrate that *C. difficile* receives the largest number and magnitude of incoming negative interactions. We identify molecular mechanisms of inhibition including acidification of the environment and competition over glucose. We demonstrate that *C. difficile*’s close relative *Clostridium hiranonis* strongly inhibits *C. difficile* via a pH-independent mechanism. While increasing the initial density of *C. difficile* can increase its abundance in the assembled community, the community context determines the maximum achievable *C. difficile* abundance. Our work suggests that the *C. difficile* inhibitory potential of defined bacterial therapeutics can be optimized by designing communities that feature a combination of mechanisms including species richness, environment acidification, and resource competition.

## Introduction

Interaction with native members of human gut microbiota inhibits the ability of gastrointestinal pathogenic strains of *Clostridioides difficile*, *Salmonella enterica* and *Escherichia coli* to secure an ecological niche and cause infection^1^. The importance of colonization resistance by gut microbiota has been particularly highlighted in *C. difficile* infections, where treatment with fecal microbiota transplants (FMT) from healthy donors has proven astonishingly effective in eliminating the symptoms of *C. difficile*^2^. Because FMT has notable risks including the transfer of antibiotic resistant organisms, potential associations with flares of inflammatory bowel disease, and in rare cases death^3–5^, defined bacterial therapeutics that have been well-characterized and standardized are needed to improve the safety and reproducibility of living bacterial therapeutic treatments. However, a key challenge to the design of effective and safe bacterial therapeutics is the vast design space of presence and absence of hundreds to thousands of potential organisms. Improving our understanding of the ecological principles of community resistance to *C. difficile* invasion could guide the design of maximally effective and safe therapeutics.

Multiple synthetic communities that inhibit *C. difficile* either *in vitro* or *in vivo* using murine models have been identified^6–11^. The majority of the defined communities are found by screening reduced complexity communities composed of isolates from a stool sample. The isolates are combined either randomly or selected based on phylogenetic diversity^6,7,10^. Other *C. difficile* inhibiting communities have been more rationally designed based on predicted mechanisms of resource competition^8^ or statistical analyses of human and murine gut microbiome data that identify taxa that correlate with infection resistance^9^. However, the design process for therapeutic synthetic microbial communities frequently does not exploit quantitative information of inter-species interactions or molecular mechanisms. A deeper understanding of the ecological principles of communities that inhibit *C. difficile* could inform the rational design of therapeutic consortia.

In macroecology, there is a long history investigating principles of invasion that has been more recently applied to microbial systems^12^. Invasion theory has identified four fundamental processes that determine the outcome of an invasion: dispersal, selection, drift, and diversification^13^. Biotic selection has been shown to be a key determinant of the outcome of an invasion, wherein higher diversity communities can competitively exclude an invader by reducing the availability of ecological niches and efficiently utilizing resources^14–16^. However, community biodiversity does not always correlate with invasion outcome, as other biotic interactions (*e.g.,* production of antimicrobial molecules), abiotic selection factors (*e.g.,* environmental pH, resource availability) and factors from dispersal, drift, and diversification processes each contribute to the outcome of invasion. For instance, in the case of a plant pathogen, the structure of the resource competition network was a better predictor of invasion outcome than biodiversity^17^. In multiple invasions of microbial communities, the dispersal factor of the initial invader abundance (i.e. propagule pressure), was found to be the key determinant of the outcome of invasion^16,18,19^.

Synthetic communities composed of known organisms can be used to investigate the driving factors of invasion outcome^16,17^. Synthetic communities enable control of initial inoculum (i.e., organism presence/absence and initial abundance), which can be manipulated to understand the ecological and molecular mechanisms influencing invader growth. Dynamic computational models informed by the experimental measurements such as the generalized Lotka-Volterra (gLV) model can be used to decipher microbial interactions and predict community assembly^20–22^. Previous modeling efforts with synthetic communities have revealed that pairwise interactions are informative of community assembly, making this approach a powerful way to understand multi-species communities with a reduced number of experiments^23^.

In this work, we use a defined synthetic gut community that represents the phylogenetic diversity of natural gut microbiota to study how principles of invasion theory apply to *C. difficile* invasion of gut microbiomes. To decipher microbial interactions and make predictions of community assembly and invasion, we use our data to construct a gLV model of our system and demonstrate that our model can accurately predict community assembly. Based on the inferred gLV interaction network, we demonstrate that negative interactions dominate the growth of *C. difficile*, which is a unique feature compared to all other species in our system. To investigate the ecological factors influencing invasion, we study the effect of propagule pressure and species richness on *C. difficile* growth. Our results show that species richness and *C. difficile* abundance exhibit a strong negative relationship across a wide range of community contexts. While increasing the propagule pressure of *C. difficile* can increase its abundance up to a maximum threshold, this threshold is dictated by the microbial community context and the ecological network. By characterizing a set of low richness communities that exhibit a range of *C. difficile* abundances, we identify multiple mechanisms that contribute to the inhibition of *C. difficile* growth including resource competition and external pH modification, highlighting that the mechanisms of inhibition of *C. difficile* vary across community contexts. Lastly, we identify a key closely related species, *Clostridium hiranonis,* that inhibits *C. difficile* growth in different synthetic communities. Our data show that microbial communities feature a wide range of resistances to *C. difficile* and multiple mechanisms of *C. difficile* inhibition, which motivates exploiting information about ecological and molecular mechanisms to design bacterial therapeutics to inhibit *C. difficile*.

## Results

### *C. difficile* coexists in co-culture with a majority of a selected set of gut microbes

We sought to understand the ecological principles of *C. difficile* invasion using synthetic gut communities (**Fig. 1a**). As a representative community, we chose a consortium of 13 prevalent gut microbes spanning the major human gut phyla *Bacteroidetes*, *Firmicutes*, *Actinobacteria, and Proteobacteria*^24^. The community features *Clostridium scindens*, a species previously shown to inhibit growth of *C. difficile* in gnotobiotic mice^9^, and a well-characterized set of 12 diverse species whose interactions on community assembly have been previously studied and computationally modeled^23^ (**Fig. 1b**).

**Figure 1:**
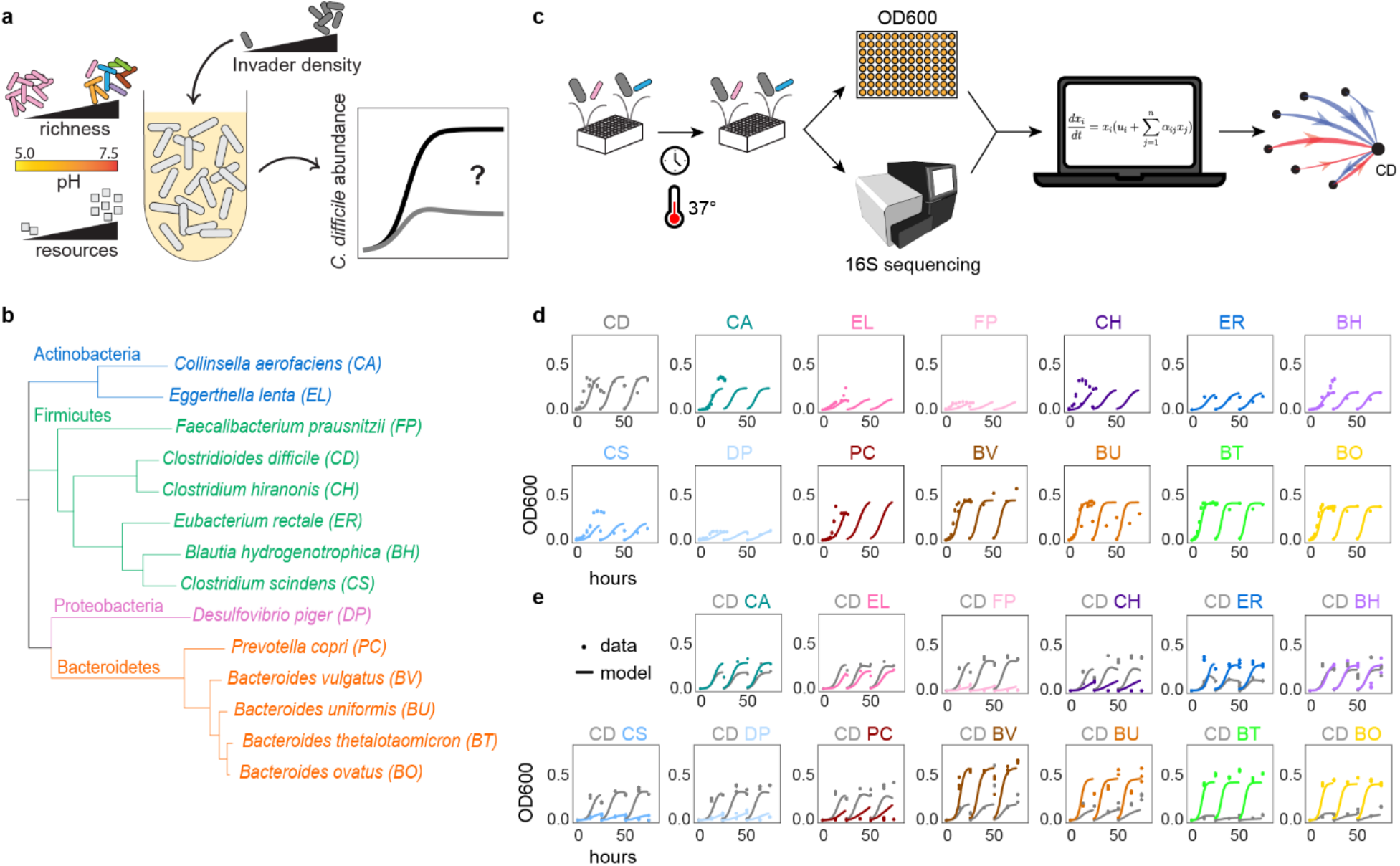
Investigating the ecological principles of *C. difficile* invasion using a diverse synthetic human gut community. **(a)***C. difficile* (CD) invasibility is hypothesized to depend on initial invader density, species richness, environmental pH, and resource availability. **(b)** Phylogenetic tree of 13-member resident synthetic gut community and *C. difficile* based on concatenated alignment of 37 marker genes. **(c)** Schematic of experimental and modeling workflow. Synthetic communities are cultured in microtiter plates in anaerobic conditions and incubated at 37°C. The absolute abundance of each species is determined by measuring cell density at 600nm (OD600) and community composition using multiplexed 16S rRNA sequencing. Absolute abundance data is used to infer the parameters of a generalized Lotka-Volterra (gLV) model. **(d)** Absolute abundance (OD600) of monospecies over time for three growth cycles. Datapoints indicate experimental biological replicates. Lines indicate simulations using the generalized Lotka-Volterra Full Model. **(e)** Absolute abundance (OD600) of *C. difficile* pairs over time for three growth cycles. First growth cycle inoculated at an equal abundance ratio of *C. difficile* to resident species based on OD600 measurements. Datapoints indicate experimental data replicates. Lines indicate simulations using the generalized Lotka-Volterra Full Model.

We used this synthetic gut community to investigate inter-species interactions influencing *C. difficile* growth. To decipher inter-species interactions driving *C. difficile* growth, we assembled combinations of species in microtiter plates in an anaerobic chamber and measured cell density by absorbance at 600 nm (OD600) and community composition by 16S rRNA gene sequencing at timepoints of interest (**Methods**). Time series measurements of species absolute abundance were used to infer the parameters of the gLV model to analyze and predict the growth dynamics of communities and deduce inter-species interactions (**Fig. 1c**). The gLV model is a system of coupled ordinary differential equations that captures the growth rate and intra-species interactions of single species and inter-species interactions that modify the growth dynamics of each species. The gLV model can be used to decipher inter-species interactions and predict the dynamics of all possible sub-communities within a larger system^23,25^ and thus can be used to study the inter-species interactions between *C. difficile* and the resident gut community (i.e. all species excluding *C. difficile*).

We first characterized the temporal behavior of pairwise communities of *C. difficile* with each resident gut bacteria since we hypothesized that these direct interactions would have the largest impact on *C. difficile* growth compared to the interactions between resident gut bacteria. To this end, each resident species was grown alone and in co-culture with *C. difficile*, specifically the R20291 reference strain of the epidemic ribotype 027 (**Fig. 1d,e**). Since variation in initial species proportions have been shown to influence community assembly^23,26^, we inoculated the pairs at 1:1 and 1:9 ratios of *C. difficile* to resident species based on OD600 values (**Fig. 1e, Fig. S1**). The communities were serially transferred every 26 hours to observe community assembly over multiple batch culture growth cycles to understand the longer-term behavior of the consortia.

*C. difficile* and the resident species coexisted (both species present at greater than 0.05 OD600 after 24 hours) in 25 of 33 (76%) conditions of 1:1 initial ratio, and 23 of 33 (70%) conditions of 1:9 initial ratio (**Fig. 1e, Fig. S1**). This frequency of coexistence in pairwise consortia was similar to a previous study that characterized the 12 member resident community, wherein 1:1 initial ratios resulted in 72% coexistence and 5:95 initial ratio resulted in 60% of pairs coexisting^23^. In both cases, equal initial ratios yielded higher rates of coexistence, consistent with the observations that initial conditions are important determinants of community assembly.

Although *C. difficile* and Bacteroides species co-existed in co-culture, the growth of *C. difficile* was reduced compared to its monospecies growth. *Bacteroides thetaiotaomicron* and *Bacteroides ovatus* strongly inhibited *C. difficile* growth, reducing *C. difficile*’s maximum carrying capacity in the first growth passage to 8% and 23% of its monospecies carrying capacity, while *Bacteroides uniformis* and *Bacteroides vulgatus* moderately inhibited *C. difficile*’s carrying capacity to 50% and 64% of its monospecies carrying capacity (**Fig. 1d,e**). Bacteroides species have been shown to inhibit *C. difficile* growth^8,10,27^ via suggested mechanisms of competition for mucosal carbohydrates or toxicity due to secondary bile acids^8,27^. Because our media does not contain mucins or bile acids, the observed inhibition indicates a separate inhibition mechanism of *C. difficile* by *Bacteroides* species. We also identified closely related species that inhibit *C. difficile* including *C. hiranonis*, the closest relative to *C. difficile* in the system (**Fig. 1b**), which reduced *C. difficile* maximum carrying capacity in the first growth passage to 78% of its monospecies carrying capacity, and the next closest relative *Eubacterium rectale*, which reduced *C. difficile’s* carrying capacity to 40% of its monospecies carrying capacity (**Fig. 1d,e**).

### *C. difficile* abundance in multispecies communities depends on species richness

We next sought to understand if the growth inhibitions of *C. difficile* observed in the majority of pairwise communities persisted in multispecies communities and to investigate the ecological principles governing *C. difficile*’s growth in multispecies communities. We designed a set of 2-13 member resident communities to experimentally characterize based on a model trained on our monospecies data (**Fig. 1d**),pairs data **(Fig. 1e)**, and previously published data of resident species pairs^23^. We inferred an initial set of parameters of the gLV model (“Preliminary Model”, **Fig. S2a, Table S3**) based on these data (**Table 1**) and used the model to predict the abundance of *C. difficile* at 48 hours in all possible 2-13 member resident communities (8,178 total communities, **Fig. S2b**). Using the predictions from the Preliminary Model, we selected a set of 94 communities whose *C. difficile* abundance at 48 hours spanned the full range of predicted *C. difficile* abundances and featured approximately equal representation of species at various initial species richness (number of species in the resident community). We experimentally assembled these communities with an initial equal abundance of all species and measured the composition of communities after 48 hours, the time by which the majority of communities had reached a steady-state according to the Preliminary Model predictions.

**Table 1:** Data used for gLV models.

| Model | Data | Figures showing data |
| --- | --- | --- |
| Preliminary Model | Monospecies<br>Pairwise communities<br>Pairwise communities from Venturelli et al<sup>23</sup> | **1d**<br>**1e, S1** |
| Full model | Monospecies<br>Pairwise communities<br>2-13 member resident communities | **1d**<br>**1e, S1**<br>**2a** (also shown in **2e, 3a**), **3b** (also shown in **3a, S6b, 3c**), **4a** (also shown in **4b, 4d**), **5b** |

We first looked at the relationship between initial species richness and *C. difficile* abundance. The biodiversity-invasibility hypothesis holds that species-rich communities have a higher fraction of ecological niches occupied, which reduces the availability of niches for invader species and thus enhances resistance to invasion relative to low-richness communities^28^. In agreement with the ecological theory, the final abundance of *C. difficile* decreased as a function of species richness (**Fig. 2a**). The negative relationship between species richness and *C. difficile* abundance remained the same whether richness was evaluated at the initial or final time point (**Fig. 2a, S3a**). Notably, *C. difficile* did not establish in any communities with richness greater than eight. The full community (13 resident members) excluded *C. difficile* from the community by 48 hours. This resistance of the full community was observed not only with the ribotype 027 strain, but also to three individual clinical isolates of *C. difficile* that originated from patients within 72 hours of their *Clostridioides difficile* Infection (CDI) diagnosis^29^ (**Fig. S3b, Methods**).

**Figure 2:**
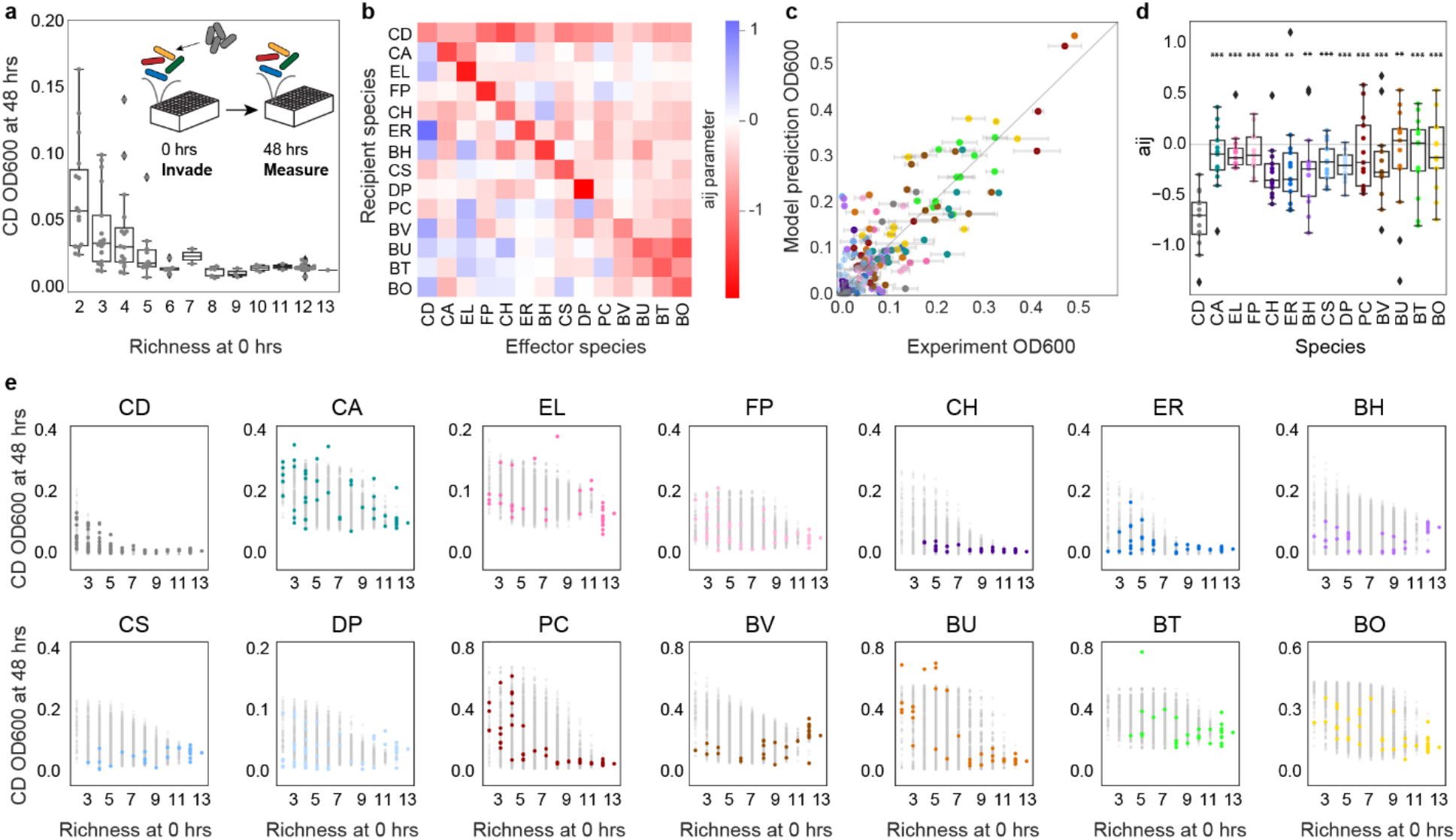
Growth of *C. difficile* decreases with community richness. **(a)** Swarmplot of *C. difficile* (CD) absolute abundance (OD600) at 48 hours in 94 sub-communities as a function of initial species richness. Datapoints indicate mean of two to three biological replicates. Line represents median, box edges represent first and third quartiles, and whiskers indicate the minimum and maximum. Outliers are denoted by diamonds. **(b)** Heatmap of inter-species interaction coefficients of the generalized Lotka-Volterra model (gLV) Full Model. **(c)** Scatterplot of average experimental absolute abundance (OD600) versus predicted species absolute abundance by the gLV Full Model in 24 held-out communities (Pearson r=0.84, p<0.001). Error bars represent standard deviation of two to three biological replicates. Gray line indicates y=x, or 100% prediction accuracy. **(d)** Box plot of incoming inter-species interactions for each species in gLV Full Model. Stars represent statistical significance between *C. difficile* and each resident species: * p<0.05, ** p<0.01, *** p<0.001 according to an unpaired t-test. Line represents median, box edges represent first and third quartiles, and whiskers indicate the minimum and maximum. Outliers are denoted by diamonds. **(e)** Subplot of the absolute abundance (OD600) of each species at 48 hours as a function of initial species richness in all 16,370 possible sub-communities of 2-13 species simulated by the gLV Full Model (gray data points) and in 94 experimentally determined subcommunities (mean-value of two to three biological replicates, colored data points).

We wanted to understand whether the strong inverse relationship between species richness and abundance was unique to *C. difficile* or also present for other species in our community. To investigate this question, we inferred a new set of gLV model parameters (“Full Model”, **Fig. 2b, Table S4**) using measurements of monospecies, pairwise and multi-species consortia (**Table 1**) and found that the Full Model had a high goodness of fit to the training data (**Fig. S4a**, Pearson r=0.89, p<0.001, model fits for monospecies and pairwise data shown in **Fig. 1d,e**). To validate the predictive capability of the Full Model, we held out 24 randomly sampled communities from the training data set that spanned a broad range of species richness and *C. difficile* abundance (**Fig. S4b**) and found that the model predicted the community composition of the held-out dataset with high accuracy (**Fig. 2c**, Pearson r=0.84, p<0.001). In contrast, the Preliminary Model trained on monospecies and pairs was substantially less predictive of these 24 multispecies communities, indicating that the model required information from the multi-species experiments (**Fig. S4c**, Pearson r=0.52, p<0.001). We performed parameter uncertainty analysis to determine if the parameters were sufficiently constrained by the data using the Metropolis–Hastings Markov chain Monte Carlo (MCMC) method (**Methods**). The coefficient of variation (CV) of the parameters ranged from 0.006 to 0.06 and 82% of parameters had a CV less than 0.05 (**Fig. S4d**), indicating that the parameters were sufficiently constrained by the data.

The Full Model’s accurate prediction of the held-out dataset indicates that the Full Model could be used to understand our system and to reliably predict multi-species community composition. Therefore, we used the Full Model to simulate the abundance of each species in all possible communities (16,383 total communities) to analyze the relationship between initial species richness and species abundance at 48 hours for all species (**Fig. 2e**). In the simulations, the 48 hour abundance of *C. difficile* displays a stronger dependence on species richness than any other species in our system (**Fig. 2e,** gray points), as evidenced by an abrupt decrease in *C. difficile* abundance for communities with greater than six species. This strong inhibition of *C. difficile* as a function of species richness can be explained by the inferred inter-species interaction network, wherein *C. difficile* displayed the largest number and magnitude of incoming negative interactions in the system (**Fig. 2d**). In addition, *C. difficile* positively impacted the growth of the majority of species in the community, which combine with the negative incoming interactions to generate a negative feedback loop on the growth of *C. difficile*. While the abundance of *C. hiranonis* and *Prevotella copri* in the subset of experimentally measured communities also exhibited a strong negative relationship with species richness (**Fig. 2e,** colored points), this trend was not observed in the model predictions of all possible communities (**Fig. 2e**, gray points). The experimentally measured communities were biased in that all communities contained *C. difficile*, such that stronger inhibition observed in the experimental set suggests *C. difficile* inhibited the growth of *C. hiranonis* and *P. copri*. This hypothesis is supported by the Full Model which features negative interactions from *C. difficile* to *C. hiranonis* and *P. copri* (**Fig. 2b**). Overall, our model analysis shows that in this system, the abundance of *C. difficile* is uniquely dependent on species richness due to a disproportionate number of negative incoming and outgoing positive inter-species interactions, leading to multiple negative feedback loops on the growth of the *C. difficile*.

### Initial abundance is a key determinant of *C. difficile* growth in synthetic communities

The propagule-pressure hypothesis dictates that increasing propagule pressure, or the amount of invader (a product of its dispersal frequency and abundance), increases the chance of a successful invasion^30^. Therefore, we next looked at the relationship between the propagule pressure of *C. difficile* and its abundance at 48 hours. In our system, we add *C. difficile* to the system a single timepoint, so the propagule pressure of *C. difficile* is equal to its initial abundance. In agreement with the theory, we found that the final abundance of *C. difficile* correlates with the initial fraction of *C. difficile* in the community (**Fig. 3a**, Pearson r=0.75, p<0.001). We analyzed the 2-13 member resident communities from our richness experiment (gray data points, **Fig. 3a**) in addition to measurements of 15, 3-4 member resident communities (**Table S2**) that we inoculated at multiple species ratios (colored data points, **Fig. 3a**). We focused on 3-4 member communities because communities in this richness range feature a wide range of *C. difficile* abundance at 48 hours (**Fig. 2a**). The 15 communities were selected to span a wide range of predicted *C. difficile* abundances and to contain communities with inferred interaction networks dominated by negative interactions, positive interactions, or approximately equal positive and negative interactions as predicted by the Preliminary Model. In all 15 communities, the abundance of *C. difficile* at 48 hours was higher in communities inoculated with a high initial density of *C. difficile* (approximately 65% of total community biomass) compared to a low initial density of *C. difficile* (approximately 10% of total community biomass) (**Fig. 3a,** inset). For five of these communities, we tested eight initial *C. difficile* densities and observed an increasing saturating function of *C. difficile* absolute abundance at 48 hours with increasing propagule pressure (**Fig. 3b**). These results demonstrate that increasing the propagule pressure of *C. difficile* can lead to higher *C. difficile* abundance in the assembled community within a given range, but beyond a threshold of initial abundance, the maximum abundance of *C. difficile* was dictated by the community context.

**Figure 3:**
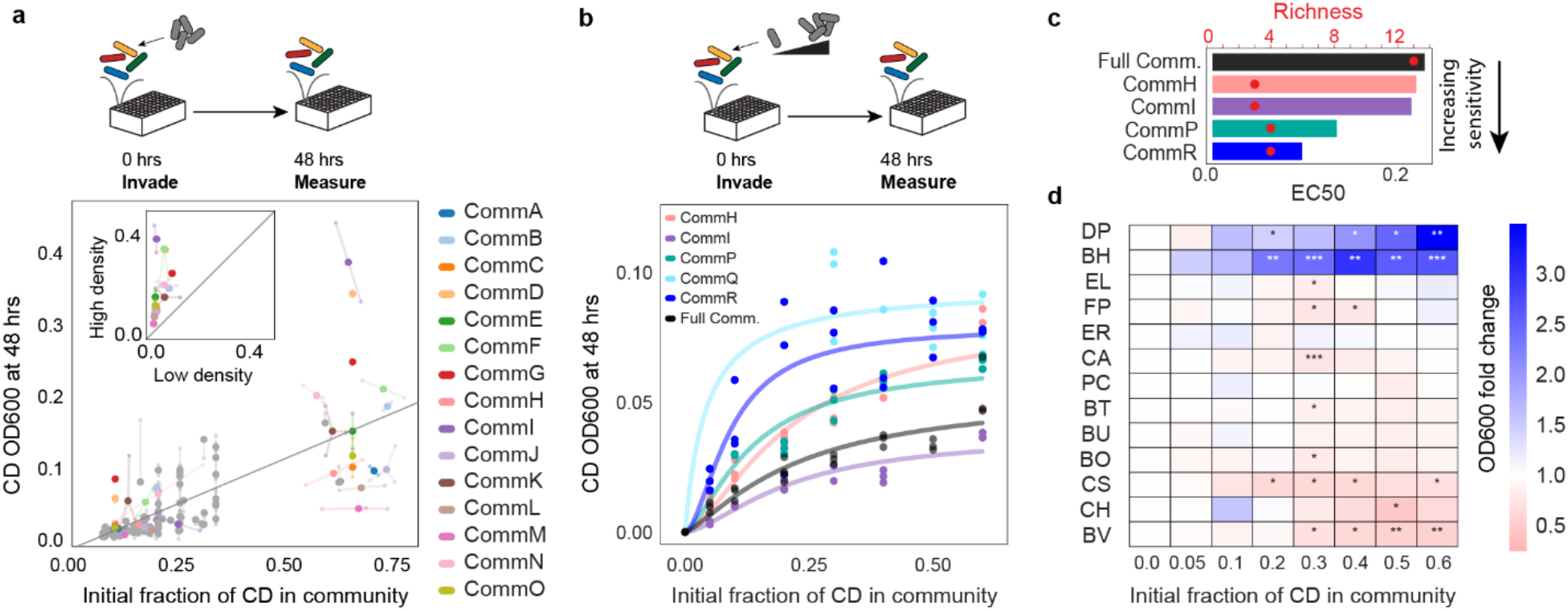
Impact of initial density on the growth of *C. difficile*. **(a)** Scatterplot of *C. difficile* (CD) absolute abundance (OD600) at 48 hours in communities as a function of the initial fraction of *C. difficile*. *C. difficile* was introduced into the communities at zero hours. Gray data points are 2-13 member resident communities measured in Fig 2a. Colored data points are 3-4 member communities measured at two initial conditions: low density (approximately 10% of total community OD600) or high density (approximately 65% total community OD600). Gray line indicates a linear regression (y=0.25x-0.01, Pearson r=0.75, p<0.001). Transparent data points indicate biological replicates and are connected to the corresponding mean values by transparent lines. Inset: Abundance of *C. difficile* at 48 hours in communities invaded with low density or high density. Gray y=x line indicates no change in abundance. Transparent data points indicate biological replicates and are connected to the corresponding mean values by transparent lines. **(b)** Absolute abundance (ODO600) of *C. difficile* at 48 hours as a function of the initial fraction of *C. difficile* in different synthetic communities. *C. difficile* was added to communities at zero hours. Datapoints indicate biological replicates. Lines indicate Hill model fits (**Methods**). **(c)** Initial fraction of *C. difficile* corresponding to the half-maximum abundance (EC50) inferred based on the fitted Hill equations in **b** for a subset of communities with sufficient measurements to constrain the function parameters. Red circles indicate the resident species richness at zero hours. **(d)** Heatmap of the fold change of species absolute abundance (mean-value of three biological replicates) in the full community with 5-60% initial *C. difficile* compared to 0% initial *C. difficile* condition. Stars represent statistical significance: * p<0.05, ** p<0.01, *** p<0.001 according to an unpaired t-test.

In the experiments and simulations, the total initial OD600 was held constant, resulting in lower initial OD600 of each species with increasing richness (**Methods**). Therefore, in light of *C. difficile*’s dependence on propagule pressure, we considered the possibility that *C. difficile*’s dependence on species richness (**Fig. 2a**) could be a result of lower initial abundance in higher richness communities. To test this possibility, we introduced a range of initial densities of *C. difficile* into the full community (richness of 13). We observed that *C. difficile* grew to a higher abundance in the full community when propagule pressure was increased, although the maximum abundance was lower than in the majority of 2-4 member communities (**Fig. 2a, 3b)**. This result indicates that while increasing propagule pressure of *C. difficile* can partially overcome the inhibiting effect of species richness, richness still decreases the maximum saturating *C. difficile* abundance.

To quantify the differential responses of the communities to varying initial *C. difficile* abundance, we defined the sensitivity to propagule pressure as the initial invader fraction that resulted in the half-maximal abundance of the invader at 48 hours, analogous to the EC50 of a dose response curve (**Fig. 3c**). The communities displayed different sensitivities to initial *C. difficile* abundance, with the EC50 ranging from 0.1 to 0.2 initial fraction of *C. difficile*. Community N (**Table S2**) was the most sensitive to invasion by *C. difficile* while the full community displayed the lowest sensitivity.

We next wanted to learn if the relationship between *C. difficile* abundance and propagule pressure changed over time. To do so, we used the Full Model to simulate *C. difficile’*s abundance in the full community from 0 to 96 hours at various propagule pressures. The simulations demonstrate that *C. difficile*’s abundance exhibits a strong dependence on propagule pressure at early times (10-20 hours), but by steady state (>48 hours) the effect of propagule pressure on *C. difficile* abundance is reduced (**Fig. S5**). The insights from the model suggest that while propagule pressure may have a significant effect on *C. difficile*’s abundance in the short term, the abundance of *C. difficile* at steady-state is dominated by other factors such as species richness and inter-species interactions.

While species richness and community composition influence *C. difficile*’s growth, we also observed that *C. difficile* had an impact on the resident community. When adding increasing amounts of *C. difficile* to six resident communities (**Fig. 3b**), we found that the composition of the resident communities at 48 hours varied as a function of the initial *C. difficile* abundance. To quantify this variation, we computed the normalized Euclidean distance between the community composition in the presence and absence of *C. difficile* (**Methods**). The Euclidean distance correlated with the abundance of *C. difficile* in the community (**Fig. S6a**, Pearson’s r=0.58, p<0.001). Mirroring our experimental data, the abundance of *C. difficile* at 48 hr correlated with the Euclidean distance between the resident community structure and the uninvaded resident community in simulations of 1-13 member resident communities invaded with *C. difficile* six hours after inoculation (**Fig. S6b**, Pearson’s r=0.61, p<0.001). Together, the experimental data and model simulations indicate that higher abundance of *C. difficile* results in a larger impact on the composition of the resident community.

In the full community, we observed that the abundance of *D. piger* and *B. hydrogenotrophica* significantly increased in communities with higher *C. difficile*, while the abundance of *B. vulgatus* significantly decreased (**Fig. 3c**). Notably, these trends were observed in the full community with the ribotype 027 strain of *C. difficile* as well as the full community with three clinical isolates of *C. difficile* (**Fig. S7a**). The interaction network from our model (**Fig. 2b**) features a positive interaction between *C. difficile* and *B. hydrogenotrophica*, suggesting that increasing initial *C. difficile* abundance directly promotes the growth of *B. hydrogenotrophica*. However, the inter-species interaction coefficients impacting *D. piger* and *B. vulgatus* were not consistent with the observed trends with these two species. These data suggest that the gLV model may not capture the effects of high initial *C. difficile* density on the growth of all resident gut species. While at high initial densities *C. difficile* significantly increased the abundance of *B. hydrogenotrophica* in the full community (**Fig. 3c**), *B. hydrogenotrophica* abundance was not affected in the 3-member communities F, G, and N (**Fig. S7b**), highlighting that *C. difficile*’s impact on a given species depends on the community context and its initial abundance. We note that *B. hydrogenotrophica* and *D. piger* share a similar metabolic niche as hydrogen consumers^31,32^, suggesting *C. difficile* could enhance their growth through a shared mechanism.

### Environmental pH is a major factor influencing *C. difficile* growth in synthetic communities

While the community experiments revealed the importance of species richness and propagule pressure on the establishment of *C. difficile* in multispecies communities, there remains unexplained variation in the data. For example, communities with the same richness invaded with equal abundances of *C. difficile* showed a wide range of *C. difficile* abundances at 48 hours (**Fig. 2a**). Since environmental pH has been shown to influence *C. difficile*’s growth in previous studies^33,34^, we turned next to investigate how biotic modification of the environment alters the growth of *C. difficile*. To this end, we grew the set of 15, 3-4 member communities for six hours and then invaded with low or high initial densities of *C. difficile*. At the time of invasion, we measured the composition of the resident community and the pH of the media (**Fig. 4a**). We also invaded the communities at zero hours with low or high initial densities of *C. difficile* to understand the role of invasion timing on the growth of *C. difficile*. *C. difficile*’s ability to establish in multiple communities significantly depended on the timing of introduction (**Fig. 4b**), indicating that biotic modification of the environment during those six hours altered *C. difficile*’s ability to grow.

**Figure 4:**
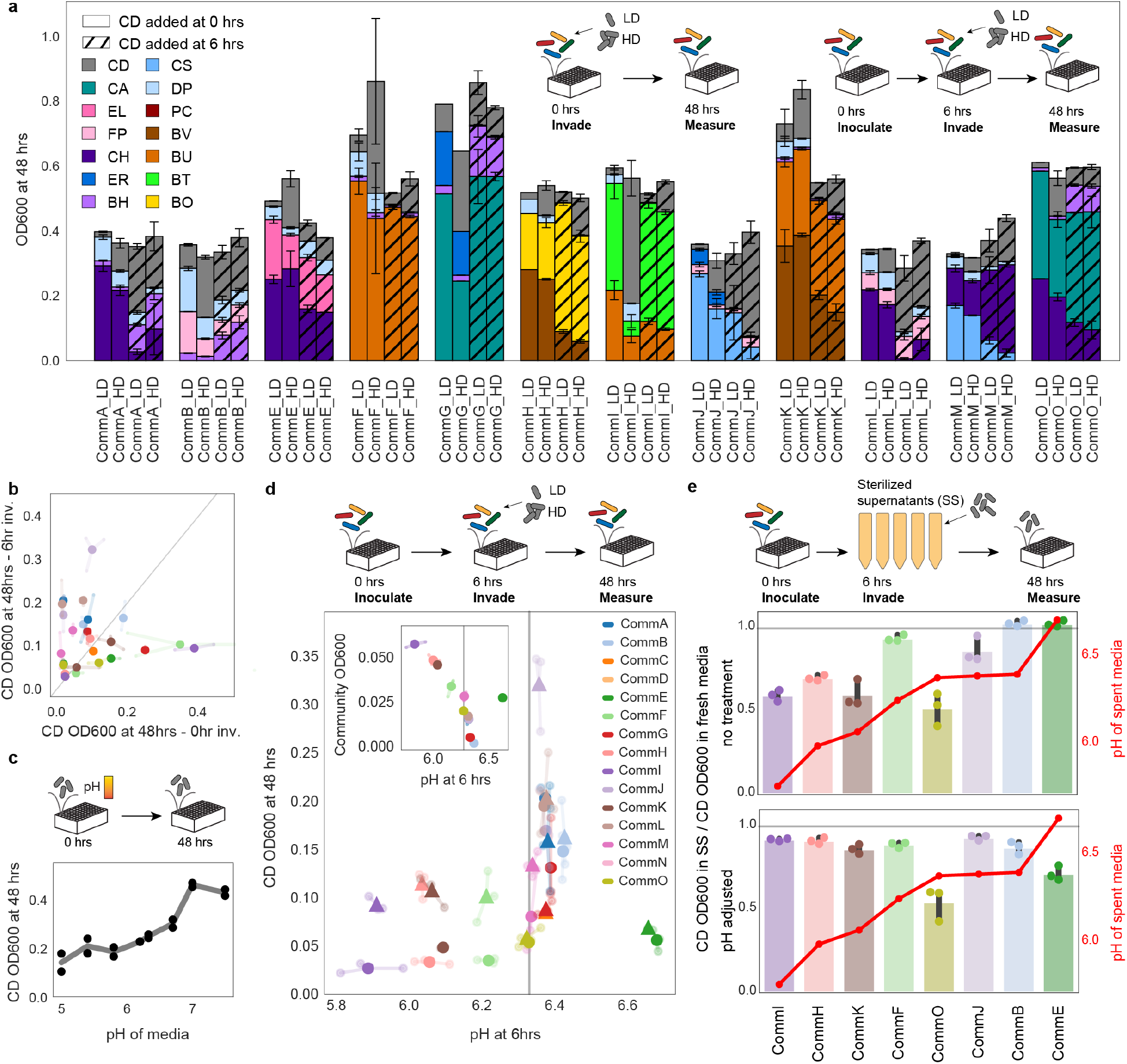
Impact of environmental factors on *C. difficile* invasion. **(a)** Barplot of composition of communities invaded by *C. difficile* at low density (“LD”) or high density (“HD”). Color indicates species identity. Hash indicates invasion time. Error bars represent one standard deviation from the mean of two to three biological replicates. **(b)** Scatterplot of the absolute abundance (OD600) of *C. difficile* at 48 hours in communities when introduced at zero hours versus six hours at low density (approximately 10% community OD600). Transparent data points indicate biological replicates and are connected to the corresponding mean values by transparent lines. Line denotes the x=y line corresponding to no change in growth. Color indicates community, see legend in **d**. **(c)** Lineplot of *C. difficile* OD600 at 48 hours as a function of the initial environmental pH. Datapoints indicate biological replicates and line indicates mean value. **(d)** Scatterplot of the absolute abundance (OD600) of *C. difficile* at 48 hours in invaded communities as a function of the environmental pH at time of invasion. Fifteen 3-4 member communities were invaded with (▲) high density *C. difficile* (approximately 33% community OD600) or (⚫) low density *C. difficile* (approximately 10% community OD600) at six hours. Color indicates community. Vertical gray line indicates pH of fresh media. Inset: Scatterplot of environmental pH and total community OD600 at six hours. Transparent data points indicate biological replicates and are connected to the corresponding mean values by transparent lines. Vertical gray line indicates environmental pH of fresh media. **(e)** Bar plot of fold change of *C. difficile* growth in sterilized supernatants (top) or supernatants where the pH was adjusted to the pH of fresh media (bottom) compared to the growth of *C. difficile* in fresh media. Growth was quantified as integral of OD600 from 0 to 20 hours. Datapoints indicate biological replicates and bars indicate mean value. Red line shows pH of community supernatants collected at six hours. Horizontal gray line indicates no change in growth compared to fresh media.

Communities that lowered the pH of the media during the first six hours featured lower *C. difficile* abundance (**Fig. 4d**). However, communities with lower pH at the time of invasion also had higher total biomass (**Fig. 4d**, inset). Since these variables are related due to growth-coupled production of acidic fermentation end products, pH or resource competition could be responsible for inhibition of *C. difficile*. Because *C. difficile* abundance increases with environmental pH (**Fig. 4c**), we hypothesized that the pH of the media contributed to growth inhibition. To confirm the contribution of pH, we grew eight of the communities harvested and sterilized the community supernatants after six hours. We grew *C. difficile* in either the filtered supernatant or a modified filtered supernatant wherein the pH was adjusted to the pH of the fresh media to eliminate the impact of pH on growth (**Fig. 4e**). For the majority of the communities, the growth phenotype of *C. difficile* in the filtered community supernatants (**Fig. 4e**) matched the growth phenotype of *C. difficile* grown in the communities (**Fig. 4d)**. However, Communities E and F inhibited *C. difficile* growth in co-culture, while the supernatants showed no significant difference in *C. difficile* growth. In Communities H, I and K, which strongly inhibit *C. difficile* in both co-culture and supernatant, increasing the supernatant pH to the pH value of fresh media eliminated the growth inhibition of *C. difficile* (**Fig. 4e**), indicating that pH was the driving factor of *C. difficile* inhibition in these community supernatants. Each of these communities contained an abundant Bacteroides species (**Table S2**) whose fermentation end products can acidify the media, suggesting abundant acidifiers are a common feature of the communities that inhibit *C. difficile*.

In contrast to this pH-dependent inhibition, the filtered supernatant of Community O (CommO) composed of *C. hiranonis, Collinsella aerofaciens* and *Blautia hydrogenotrophica*, whose pH did not significantly differ from the pH of fresh media, inhibited the growth of *C. difficile* regardless of pH adjustment (**Fig. 4e**), indicating that this community inhibits *C. difficile* via a pH-independent mechanism. *C. difficile* was not inhibited by the filtered supernatant of Community E (CommE) composed of *C. hiranonis*, *Desulfovibrio piger* and *Eggerthella lenta*, which uniquely had a higher pH than fresh media, but did inhibit *C. difficile* when the pH was reduced to the pH of fresh media (**Fig. 4e**). This suggests that the filtered supernatant promotes *C. difficile*’s growth by enhancing environmental pH and the community inhibits *C. difficile*’s growth by a separate pH-independent mechanism. The growth inhibition was only revealed when the pH increase of the media was eliminated, demonstrating an interplay of different mechanisms influencing *C. difficile* growth.

Overall, we determined that the modification of environmental pH alters *C. difficile* growth in many communities. To determine if *C. difficile*’s sensitivity to pH was unique and thus a potential mechanism contributing to *C. difficile*’s unique and strong inverse dependence on species richness (**Fig. 2e**), we measured the carrying capacity of each species as a function of environmental pH in monoculture and determined the slope of the line fit to these data (**Fig. S8a**), representing the sensitivity of species growth to external pH. Our results demonstrated that *C. difficile’*s pH sensitivity was not unique, ranking eighth most sensitive out of the 14 species (**Fig. S8b**). Therefore, while acidification of the media is one mechanism by which communities inhibit *C. difficile* in our system, our results suggest that there are also pH-independent mechanisms that contribute to a strong dependence between species richness and *C. difficile* growth.

### *C. hiranonis* inhibits *C. difficile* through a pH-independent mechanism

Notably, the two communities that displayed pH-independent growth inhibition (CommE and CommO) contained *C. hiranonis*, which has a strong bidirectional negative interaction with *C. difficile* in our Full Model (**Fig. 2b**). Our model predicted that the abundance of *C. difficile* at 48 hours decreases with increasing initial abundance of *C. hiranonis* in Communities E, O and the *C. difficile-C.hiranonis* pair. We tested this prediction experimentally and confirmed that *C. hiranonis* grew to a higher absolute abundance and *C. difficile* grew to a lower absolute abundance in communities inoculated with higher initial fraction of *C. hiranonis* (**Fig. 5b,** inset). The growth of *C. difficile* was sensitive even to low initial amounts of *C. hiranonis*, featuring a significant decrease in growth between 0% and 10% initial *C. hiranonis* in CommE (>4-fold decrease) and CommO (>1.5 fold decrease) (**Fig. 5b**). The strength of inhibition of *C. difficile* as a function of the initial density of *C. hiranonis* was substantially higher in CommE and CommO than in the *C. hiranonis*-*C. difficile* pair (**Fig. 5b**). This result indicates that the other species in the communities enhanced the inhibitory effect of *C. hiranonis* on *C. difficile* growth.

**Figure 5:**
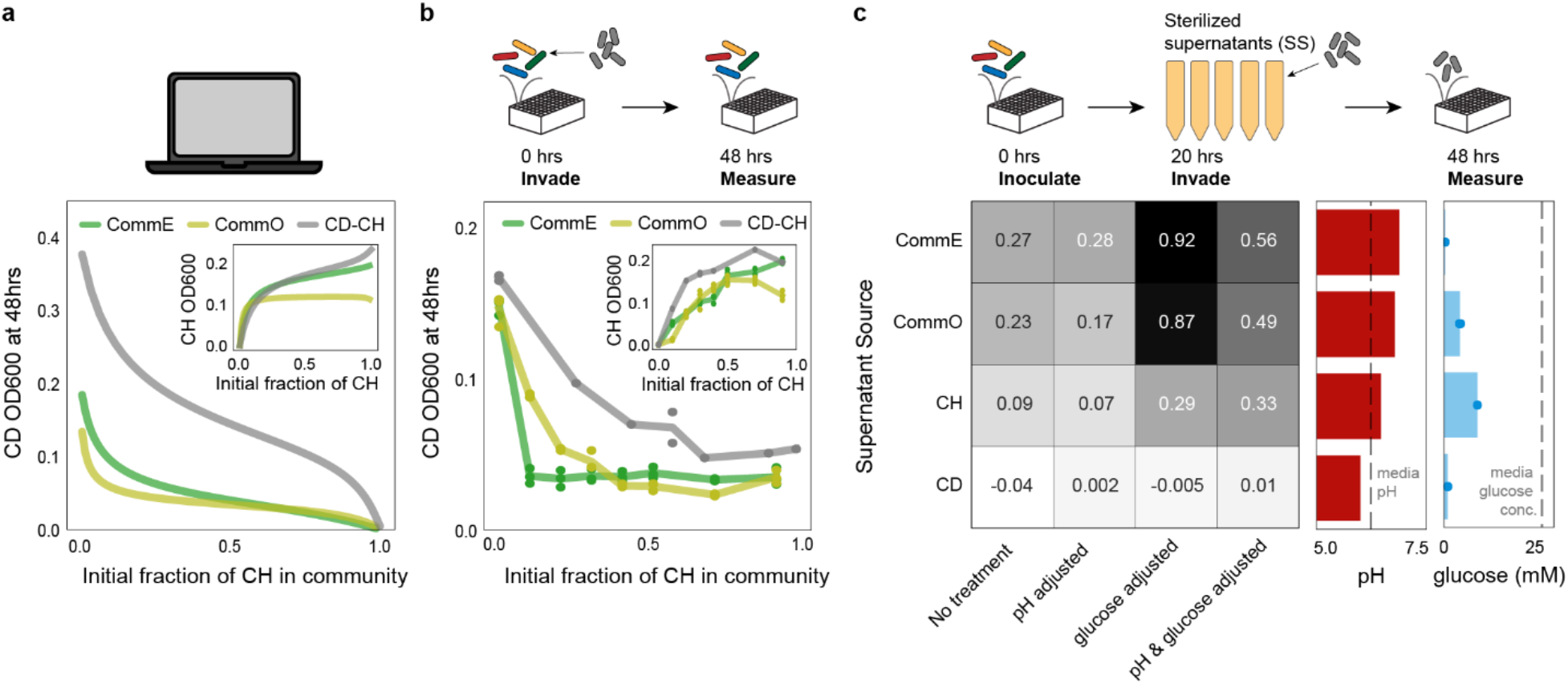
*C. hiranonis* inhibits the growth of *C. difficile*. **(a)** Lineplot of simulated *C. difficile* (CD) absolute abundance (OD600) at 48 hours using the generalized Lotka-Volterra (gLV) Full Model as a function of the initial fraction of *C. hiranonis* (CH) in different communities. Inset: Lineplot of simulated *C. hiranonis* absolute abundance (OD600) at 48 hours in the gLV Full Model as a function of initial fraction of *C. hiranonis* in the community. **(b)** Lineplot of *C. difficile* absolute abundance (OD600) at 48 hours as a function of the initial fraction of *C. hiranonis* in the community. Inset: Lineplot of *C. hiranonis* absolute abundance (OD600) at 48 hours in community as a function of initial fraction of *C. hiranonis* in the community. Datapoints indicate biological replicates and lines indicate mean values. **(c)** Heatmap of *C. difficile* growth in treated sterilized supernatants. The values of the heatmap represent the fold change between the integral of *C. difficile* OD600 from 0 to 56 hours in the treated supernatant and the integral of *C. difficile* OD600 from 0 to 56 hours in fresh media (mean-values of three biological replicates). Red barplot indicates the pH of the supernatant before treatment. Dashed line indicates the pH of fresh media. Blue barplot indicates the glucose concentration of the supernatant before treatment. Bar indicates average value and points indicate technical replicates. Dashed line indicates glucose concentration of fresh media.

We next considered the mechanism of *C. hiranonis*’s inhibition of *C. difficile*. *C. hiranonis* is known to convert primary bile acids into secondary bile acids which are inhibitory to *C. difficile*^35^, however with no primary bile acids in our media we turned to other possible inhibition mechanisms. *C. difficile* was inhibited by the filtered supernatants of CommE, CommO and *C. hiranonis* (**Fig. 4e**, **5c**), indicating the inhibition effect does not require direct cell contact, suggesting mechanisms such as production of antibiotics or toxic metabolic byproducts, competition for resources, or pH modification. In a soft agar overlay assay, where *C. difficile* grows in soft agar layered on top of a *C. hiranonis* colony, we did not see inhibition by *C. hiranonis*, although we did see zones of inhibition by specific Bacteroides species (**Fig. S9**). Because the pressures of resource competition are removed in a soft agar assay (*C. difficile* has access to resources in the soft agar layer), we hypothesized that inhibition observed in liquid culture with *C. hiranonis* was due to resource competition. The hypothesis of resource competition by *C. hiranonis* was informed by the ecological theory that closely related species are likely to compete for overlapping resource niches, which has been observed in microbial systems^36^. This theory is supported by our model which features a moderate but statistically significant positive correlation between the Full Model inferred inter-species interaction coefficients and phylogenetic distance between species (**Fig. S10,** Pearson r=0.34, p<0.001). Additionally, *C. hiranonis* has been shown to consume more metabolites than any of the other resident species in our media conditions^23^, suggesting the potential to compete with *C. difficile* over other resources.

To investigative potential mechanisms of resource competition between *C. difficile* and *C. hiranonis*, we focused on two key resources present in our media that *C. difficile* has been shown to utilize: glucose and succinate^37,38^. We measured the concentration of these resources in the supernatant of *C. difficile*, *C. hiranonis*, CommE, and CommO after 20 hours (**Methods**).

Corroborating previous data, each donor supernatant inhibited the growth of *C. difficile*, and adjusting the pH of the supernatants to the pH of fresh media did not remove the inhibition (**Fig. 4e**, **5c**). While succinate concentrations were either moderately increased or similar to the concentration in fresh media (**Fig. S11**), glucose was substantially lower in the supernatants (**Fig. 5c**). Adjusting the glucose concentration to the concentration of fresh media almost completely restored *C. difficile* growth in the CommE and CommO supernatants, but only moderately restored growth in the *C. hiranonis* supernatant (**Fig. 5c**). These results indicate that competition over glucose and not pH modification was a driving mechanism of *C. difficile* inhibition in CommE and CommO. However, neither competition over glucose nor pH modification was able to explain the inhibitory effect of *C. hiranonis* on *C. difficile* in the pairwise community, suggesting *C. hiranonis* could be inhibiting *C. difficile* by competing for a different resource. Therefore, our results suggest that there are multiple mechanisms of *C. difficile* inhibition by *C. hiranonis* and the other resident gut bacteria and that these mechanisms depend on community context.

## Discussion

We combined bottom-up construction of microbial communities with dynamic computational modeling to investigate microbial interactions impacting the growth of *C. difficile*. Our work demonstrates that microbial communities feature a wide range of resistances to *C. difficile* invasion. This variability in invasion outcome as a function of community context indicates that the choice of organisms is a major design factor that can be optimized to treat *C. difficile* infections and motivates exploiting ecological information in the design process. Previous efforts to design defined consortia for *C. difficile* inhibition used top-down selections by reducing the complexity of cultured fecal samples alone or combined with screening of antibiotic resistance phenotypes^6,7^. Some consortia have been designed by combining selected species in a bottom-up approach, but we note that these selections use a single design criterion^8,9^. Beyond previously demonstrated mechanisms of bile acid transformations^9^ and mucosal sugar competition^8^, our results demonstrate that acidification of the environment and competition over limiting resources such as glucose can inhibit *C. difficile* growth. Further, species richness was a driving factor of *C. difficile* growth across a wide range of community contexts. In sum, these results suggest that multiple mechanisms could be combined to design an optimal defined bacterial therapeutic to inhibit *C. difficile*.

Studies have shown that gut microbiomes of patients with CDI have significantly lower richness than healthy controls^39,40^, but this association does not distinguish whether CDI reduces the richness of gut microbiomes or low richness microbiomes are more susceptible to CDI. The striking trend between richness and *C. difficile* abundance in our data suggests that low richness microbiomes are more susceptible to CDI. Supporting this hypothesis, the susceptibility of low richness communities to invasion has been demonstrated in other microbial systems^14,41^. This suggests that the low gut microbiome richness induced by antibiotics^42^ could contribute to increased CDI risk after antibiotic use^43^. Additionally, the efficacy of FMTs may be due to the high richness of stool samples which are estimated to have greater than one hundred species^44^.

Based on our work, high richness communities would be the most effective bacterial therapeutics to inhibit *C. difficile* colonization. The scalable manufacturing of high richness bacterial therapeutics is challenging, indicating the need for new bacterial manufacturing techniques to reliably culture communities of gut species as opposed to single species, while maximizing evenness and growth. Nevertheless, if scalable manufacturing of high richness communities remains an unresolved challenge, our work suggests it is possible to design low richness inhibitory communities. While all high richness communities (eight species or more) excluded *C. difficile* in our system, we did find low richness communities that excluded *C. difficile*. For example, the 3-member Community I excluded *C. difficile* as effectively as the full community, featuring a similar maximum *C. difficile* abundance as a function of initial *C. difficile* density (**Fig. 3b**). Corroborating these results, low richness communities as small as 5-7 members have been shown to inhibit *C. difficile in vitro* and in murine models^7–9^.

Our results demonstrated that communities can inhibit growth of *C. difficile* by acidifying the environment. We showed that communities that reduce the external pH below 6.2 inhibit *C. difficile* in a pH-dependent manner, consistent with studies showing that *C. difficile* has lower viability and rates of sporulation in acidic environments^33,34^. While our *in vitro* system lacks the pH-buffering secretion of bicarbonate by host intestinal epithelial cells, the amount of bicarbonate buffer in our media (4.8 mM) is within the estimated range in the gastrointestinal tract (2-20mM)^45^, suggesting the observed pH changes in our media could be physiologically relevant. Even with host bicarbonate secretions that regulate the pH of the gut, fermentation by colonic bacteria impacts luminal pH, which can be manipulated using dietary substrates^46^. Notably, a human cohort study found a strong association between alkaline fecal pH and CDI^47^. Together, these suggest that manipulation of the pH of the gut environment via bacterial therapeutics or dietary interventions is a potential microbiome intervention strategy to inhibit *C. difficile*. To optimize inhibitory potential of bacterial therapeutics, in addition to designing communities that acidify the environment, communities could also be designed to maximize resource competition between resident members and *C. difficile*. We found that relieving resource competition through addition of glucose reduced *C. difficile* inhibition by 22-90% depending on the community context (**Fig. 5c**). Therefore, constituent members of the bacterial therapeutics that compete with *C. difficile* for the estimated 20% of carbohydrates, such as glucose, that escape absorption by the host^48,49^ could reduce *C. difficile* colonization.

We find that increasing the propagule pressure of *C. difficile* leads to an increase in the pathogen’s abundance in the community (**Fig. 3a,b**). While propagule pressure has been shown to determine invasion success in microbial invasions^16,18,19^, here we demonstrate that this applies to *C. difficile* in synthetic gut communities. Propagule pressure is known to be important in murine *C. difficile* infections, where mice cohoused with supershedders containing 10^8^ CFU g^−1^ *C. difficile* in their feces became colonized with *C. difficile*, whereas mice cohoused with low shedders containing 10^2^ CFU g^−1^ *C. difficile* did not become colonized^50^. However, the relationship between *C. difficile* dosage and incidence of CDI in humans is unknown. Our results suggest that the density of *C. difficile* could be an important variable in the outcome of *C. difficile* invasions in a clinical setting. In our experiments, we found that while increasing the propagule pressure of *C. difficile* increases its abundance in the community over a range of initial densities, communities varied in the maximum *C. difficile* abundance (**Fig. 3a,b**). This suggests that different human gut microbiome compositions vary in their resistance to invasion of varying amounts of *C. difficile* due to ecological interactions.

We were able to construct a gLV model that accurately predicts the composition of 2-13 member communities by training on similarly complex data (1-13 members), but parameters trained on low richness communities alone (1-2 species) were not able to predict these higher richness communities, as has been seen previously^25^. The inferred inter-species interaction network was dominated by competition, with 73% negative interactions (*α*_ij_ < −0.01), consistent with the large number of negative interactions observed in other microbial communities^25,51^. Notably, *C. difficile* was the only species that was inhibited by all other community members. Infection by *C. difficile* disrupts the environment of gut bacteria by causing diarrhea (i.e. reduces residence time for gut bacteria), inducing intestinal inflammation, and altering the resource landscape^52^, suggesting the possibility that gut bacteria have evolved to negatively impact the growth of *C. difficile* in order to promote their fitness in the gut.

Bacteroides have been found to both inhibit and promote *C. difficile* growth in different environments^8,10,38,53^, but in our system all Bacteroides species in the community strongly inhibited *C. difficile*. We did not observe a strong inhibition of *C. difficile* by *C. scindens* which has been documented to occur via production of secondary bile acids that inhibit *C. difficile* germination^9^ because our media does not contain bile acids. Instead, in our system the closest relative of *C. difficile*, *C. hiranonis*, was the strongest inhibitor of *C. difficile* abundance. Currently, phylogenetic relatedness is a major design factor used to select species for defined bacterial therapeutics. For example, defined bacterial therapeutics have been constructed by treating fecal samples with ethanol to select for spore-forming bacteria, which are primarily closely related *Clostridiales* species^54^. Our work shows that including other more diverse species in CommE and CommO resulted in stronger inhibition of *C. difficile* as a function of *C. hiranonis* initial abundance. Therefore, while our results demonstrate that closely related species can inhibit *C. difficile*, including other diverse commensal bacteria in the community could substantially increase the degree of inhibition.

In sum, we identified ecological and molecular mechanisms of resistance to invasion by *C. difficile* using a synthetic gut microbiome. While our system lacks the full diversity of the human gut microbiome and a host-interaction component, many of our results support principles of invasion theory based on a broad range of systems, suggesting that some of these principles could be generalized to the mammalian gut environment. Future work could create panels of gut microbial communities that feature different weightings of the multiple community resistance mechanisms demonstrated in this work. These panels could be tested *in vitro* for inhibition of *C. difficile* growth and promising candidates could be introduced into germ-free mouse models to evaluate their *C. difficile* inhibitory potential as bacterial therapeutics.

## Acknowledgements

Research was sponsored by the National Institutes of Health and was accomplished under Grant Number R35GM124774 and National Institute of Biomedical Imaging and Bioengineering under grant number R01EB030340. S.E.H. was supported in part by the National Institute of General Medical Sciences of the National Institutes of Health under Award Number T32GM008349. R.L.C. was supported in part by a National Human Genome Research Institute training grant to the Genomic Sciences Training Program under Award Number T32HG002760.

## Author contributions

O.S.V. and S.E.H. conceived the study. S.E.H. carried out the experiments. S.E.H. and Y.Q. performed computational modeling and analysis. R.L.C. and Y.Q. wrote customized scripts to perform modeling analyses. S.E.H. and O.S.V. analyzed and interpreted the data. O.S.V. secured funding. N.S. and L.W. isolated clinical *C. difficile* strains and provided information on these strains. S.E.H. and O.S.V. wrote the paper and all authors provided feedback on the manuscript.

## Methods

### Strain information and starter culture inoculations

Cells were cultured in an anaerobic chamber (Coy Lab products) with an atmosphere of 2.5±0.5% H_2_, 15±1% CO_2_ and balance N_2_. The strains used in this work were obtained from the sources listed in **Table S1**. The three clinical *C. difficile* isolates (MS002, MS010, MS011) were *C. difficile* NAAT (GeneXpert) positive via admission stool sample and toxin A (tcdA) and toxin B (tcdB) positive via in-house research PCR. Each patient was diagnosed with and treated for CDI. Single-use glycerol stocks were prepared as described previously^25^. Single species starter cultures were inoculated by adding 100 μL of a single-use 25% glycerol stock to 5mL of Anaerobic Basal Broth media (ABB, Oxoid). *E. rectale* starter cultures were supplemented with 33 mM Sodium Acetate (Sigma-Aldrich) and *D. piger* starter cultures were supplemented with 28 mM Sodium Lactate (Sigma-Aldrich) and 2.7 mM Magnesium Sulfate (Sigma-Aldrich). To begin experiments with organisms in similar growth phases, starter cultures were inoculated either 16 hours or 41 hours prior to experimental set up, depending on the growth rate of the organism (**Table S1**).

### Monospecies and pairs experiments

Starter cultures were diluted to 0.0022 OD600 in ABB (Tecan Infinite Pro F200). For monospecies in **Fig. 1d**, diluted cultures were added directly to 96 deep well plates for final OD600 of 0.0022. For pairs in **Fig. 1e**, diluted cultures were combined into pairs in 96 deep well plates at 1:1 or 1:10 volume ratios for final OD600 of 0.0011 or 0.00022 and 0.00198. Cultures were combined using a liquid handling robot (Tecan Evo 100). Plates were covered with gas-permeable seal (BreatheEasy) and incubated at 37°C with no shaking.

### Multispecies community experiments

Starter cultures were diluted to 0.0066 OD600. Diluted cultures were combined into communities in 96 deep well plates using a liquid handling robot (Tecan Evo). The 94 sub-communities in **Fig. 2a** were created by combining equal volumes of each diluted starter culture, so the initial OD600 of each species in the community was 0.0066 divided by the number of species. The 3-4 member *C. difficile* titration communities in **Fig. 3b** were combined such that all non-*C. difficile* species had an initial OD600 of 0.00165, and *C. difficile* had an initial OD600 of 0, 0.00026, 0.00055, 0.0012, 0.0021, 0.0033, 0.00495, and 0.0074 in the 3 member communities and 0, 0.00035, 0.00073, 0.00165, 0.0028, 0.0044, 0.0066, and 0.0099 in the 4 member communities for initial fractions 0, 0.1,0.2, 0.3, 0.4, 0.5, and 0.6 respectively. The full community in **Fig. 3b** was combined so that all non-*C. difficile* species had an initial OD600 of 0.00047, and *C. difficile* had an initial OD600 of 0, 0.00032, 0.0015, 0.0026, 0.0041, 0.0061, 0.0092 for initial fractions 0, 0.1, 0.2, 0.3, 0.4, 0.5, and 0.6 respectively. The communities in **Fig. 4a** were combined so that all non-*C. difficile* species had an initial OD600 of 0.00165, and *C. difficile* had an initial OD600 of 0.00055 (10% of community) in the low density zero hour invasion condition, 0.009 (65% of community) in the high density zero hour invasion condition, and was not introduced into the six hour invasion condition. After six hours of incubation, the community OD600 was measured and *C. difficile* was added to the six hour invasion conditions so that its OD600 was 10% (low density condition) or 33% (high density condition) of the community. The *C. hiranonis* titration communities in **Fig. 5b** were combined so that all non-*C. hiranonis* species had an initial OD600 of 0.00165, and *C. hiranonis* had an initial OD600 of 0, 0.00055, 0.00012, 0.0021, 0.0033, 0.0050, 0.012 and 0.045 for initial fractions 0, 0.1, 0.2, 0.3, 0.4, 0.5, 0.7, and 0.9 respectively. Plates were covered with gas-permeable seals (BreatheEasy) and incubated at 37°C with no shaking.

### Culture sample collection

At each timepoint, samples were mixed and aliquots were removed for sequencing and for measuring OD600. We measured OD600 of two dilutions of each sample and selected the value that was within the linear range of the instrument (Tecan Infinite Pro F200). Sequencing aliquots were spun down aerobically at 3500 rpm for 15 minutes and stored at −80°C. For timepoints with dilutions, samples were mixed and aliquots were collected for sequencing and OD600 measurements before the samples were diluted 1:20 into fresh media. Abundance of the diluted sample was calculated by dividing the undiluted measurements by the dilution factor of 20.

### pH measurements and adjustments

The pH of each community in **Fig. 4d** was measured using a phenol red assay as described previously^25^. The pH of each supernatant in **Fig. 4e, 5c** was measured using a pH probe (Mettler Toledo). The pH of each supernatant was adjusted to the pH of fresh media by adding small volumes of sterile 5M NaOH and 5M HCl.

### Supernatant experiments

Starter cultures were diluted to 0.0066 OD600. Diluted cultures were combined into communities in 96 deep well plates using a liquid handling robot (Tecan Evo). Communities were created by combining equal volumes of each species, so the final OD600 of each species in the community was 0.0066 divided by the number of species. Plates were covered with gas-permeable seal (BreatheEasy) and incubated at 37°C with no shaking. After incubation time of six hours (**Fig. 4e**) or 20 hours (**Fig. 5c**), cultures were spun down aerobically at 3500 rpm for 15 minutes and sterile filtered using Steriflip 0.2 μM filters (Millipore-Sigma) before returning to anaerobic chamber. Media controls were spun down and filtered aerobically in parallel with samples. *C. difficile* was inoculated in the sterilized supernatants to a final OD600 of 0.0022 in 96 well microplates that were covered with gas-permeable seals (BreatheEasy), incubated at 37°C with shaking, and OD600 was measured every 2 hours (Tecan Infinite Pro F200).

### Genome extractions

Genomic DNA was extracted using a method adapted from previous work^25^. Briefly, cell pellets were resuspended in 180 μL Enzymatic Lysis Buffer containing 20 mg/mL lysozyme (Sigma-Aldrich), 20 mM Tris-HCl pH 8 (Invitrogen), 2 mM EDTA (Sigma-Aldrich), and 1.2% Triton X-100 (Sigma-Aldrich). Samples were incubated at 37°C at 600 RPM for 30 minutes. Samples were treated with 25 μL 20 mg/mL Proteinase K (VWR) and 200 μL Buffer AL (Qiagen), mixed by pipette and incubated at 56°C at 600 RPM for 30 minutes. Samples were treated with 200 μL 200 proof ethanol (Koptec), mixed by pipette and transferred to 96 well nucleic acid binding plates (Pall). After washing with 500 μL Buffer AW1 and AW2 (Qiagen), a vacuum was applied for 10 minutes to dry excess ethanol. Genomic DNA was eluted with 110 μL Buffer AE (Qiagen) preheated to 56°C and then stored at −20°C.

Genomic DNA was quantified using Sybr Green fluorescence assay with a 6-point DNA standard curve (0, 0.5, 1, 2, 4, 6 ng/μL Biotium). 1 μL of samples and 5 μL of standards were diluted into 95 μL of 1X SYBR Green (Invitrogen) in TE buffer and mixed by pipette. Fluorescence was measured with an excitation/emission of 485/535 nm (Tecan Spark). Genomic DNA was normalized to 1 ng/μL in molecular grade water using a liquid handling robot (Tecan Evo 100). Samples less than 1 ng/μL were not diluted. Diluted genomic DNA was stored at −20°C.

### Primer design, library preparation, and sequencing

Dual-indexed primers for multiplexed amplicon sequencing of the 16S v3-v4 region were designed as described previously^23,25^. Briefly, oligonucleotides (Integrated DNA Technology) were arrayed into 96 well plates using an acoustic liquid handling robot (Echo LabCyte) and stored at −20°C. Genomic DNA was PCR amplified using Phusion High-Fidelity DNA Polymerase (Thermo-Fisher) for 25 cycles with 0.05 μM of each primer. Samples were pooled by plate, purified (Zymo Research), quantified by NanoDrop and combined in equal proportions into a library. The library was quantified using Qubit 1x HS Assay (Invitrogen), diluted to 4.2 nM, and loaded at 21 pM onto Illumina MiSeq platform for 300-bp paired end sequencing.

### Data Analysis

Sequencing data was analyzed using a method adapted from previous work^23^. MiSeq Reporter software demultiplexed the indices and generated FastQ files. FastQ files were analyzed using custom python scripts. Paired reads were merged using PEAR (Paired-End reAd mergeR) v0.9.0 (Zhang et al, 2014). A reference database containing 16S v3-v4 region of each species in the study was created by assembling consensus sequence based on sequencing results of each monospecies. The classify.seqs command in mothur was used to map reads to the reference database using the Wang method with a confidence cut off of 60% (Wang et al). Relative abundance was calculated by dividing the read counts mapped to each organism by the total reads in the sample. Absolute abundance was calculated by multiplying the relative abundance of an organism by the OD600 of the sample.

### Glucose and succinate quantification

Succinate concentration was quantified using EnzyChrom Succinate Assay Kit (BioAssay Systems) with two technical replicates of each filtered supernatant and ABB media diluted 1:100 in buffer to fall in the linear range of the calibration curve. Glucose concentration was quantified using Amplex Red Glucose Assay Kit (ThermoFisher) with four technical replicates of each filtered supernatant and ABB media diluted 1:100 in buffer to fall in the linear range of the calibration curve. Glucose of the supernatants was adjusted to the concentration of glucose in ABB media using a filter sterilized glucose stock (Alfa Aesar).

### Soft agar overlay

Starter cultures (3 μL) were spotted in triplicate on 1.5% ABB agar plates and incubated for 24 hours. At this time, colonies were killed via aerobic exposure for six hours and then returned to anaerobic conditions. Soft 0.7% ABB agar was inoculated to 0.0022 OD600 *C. difficile* and poured over the colonies. Plates were then incubated for 24 before analyzing and imaging zones of inhibition.

### Generalized Lotka-Volterra Model

The gLV model is a set of *N* coupled first-order ordinary differential equations:

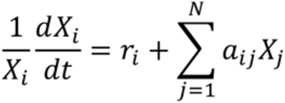

where *N* is the number of species, the parameter *X_i_* is the abundance of species *i*, the parameter *r_i_* is the basal growth rate of species *i*, the parameter *α_ij_*, called the interaction parameter, is the growth modification of species *i* by species *j* and the parameter *X_j_* is the abundance of species *j*. The parameter *α_ij_* is constrained to be negative when *i*= *j*, representing intra-species competition.

### Parameter estimation

The gLV model parameters were estimated from time-series measurements of single-species and multispecies cultures using the nonlinear programming solver FMINCON in MATLAB, which finds the optimal set of parameters that minimizes a given cost function. The estimation was implemented using previously developed custom MATLAB scripts^25^. The cost (C) of the optimization algorithm was computed by (1) simulating each species *m* in each community *k* with an ODE solver and summing the mean squared error between the abundance of each species in the simulation *X_model_* and data *X_exp_* at each timepoint *n* (2) adding the sum each parameter *θ* squared multiplied by a regularization coefficient λ:

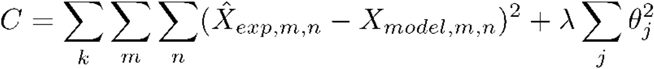

The second step is a L2 regularization, which penalizes the magnitude of the parameter vector to prevent overfitting the data. The optimization was repeated with a range of regularization coefficients. The regularization coefficient that resulted in a parameter set with a mean squared error of 110% of the non-regularized parameter set was selected, which was λ=0.5 for the Preliminary model and λ =0.1 for the Full model. The data used for parameter estimation for the Preliminary model and Full model are given in **Table 1**. To validate the predictive ability of the model, 24 2-13 member resident communities (**Fig. S4a**) were left out from the training data set and a set of parameters was inferred from this reduced data set using λ =0.1 for the regularization coefficient. The community compositions of the 24 held-out communities were simulated with this parameter set to evaluate the predictive capability of the model on held-out data (**Fig. 2c**)

### Parameter uncertainty analysis

To quantify the uncertainties in gLV parameters, an adaptive Markov Chain Monte Carlo (MCMC) method was used to sample from the posterior gLV parameter (θ) distribution *P*(θ|**y**) given a sequence of *m* abundance measurements **y**=(**y**_1_,…,**y**_m_). In particular, for the *k*-th measurement, **y**_*k*_ is a vector that concatenates all abundance measurements collected from all sub-community experiments. Uncertainty for the *k-*th measurement was modeled by an additive and independent noise, which is distributed according to 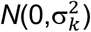, where 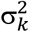 is the diagonal covariance matrix for experimental data collected in the *k-*th measurement. Given a fixed parameter θ, the gLV model was simulated to obtain the model predicted abundance 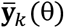 at every instant *k*. The likelihood to observe a sequence of abundance measurements **y** was then computed as:

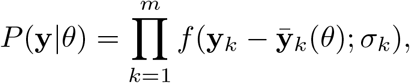

where *f*(∙; σ_k_) is the probability density function for the normal distribution 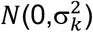. The posterior distribution was then described according to Bayes rule as *P*(θ|**y**) ∝ *P*(**y**|θ)*P*(θ), where *P*(θ) is the prior parameter distribution. Normal priors were used for the parameters. The means of the normal distributions were set to the parameters estimated by the FMINCON method and the coefficients of variation were set to 5%.

An adaptive, symmetric, random-walk Metropolis MCMC algorithm^55^ was then used to draw samples from this posterior distribution. Specifically, given the current sample θ^(*n*)^ at step *n* of the Markov chain, the proposed sample for step (*n*+1) is θ^(*n*+1)^ = θ^(*n*)^+δ^(*n*)^, where δ^(*n*)^ is drawn from a normal distribution. The algorithm is adaptive in the sense that the covariance of this normal distribution is given by 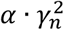, where 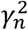 is the covariance of θ^(1)^,…, θ^(*n*)^ and ± is a positive parameter. The proposed sample is accepted with probability 1 if *P*(θ^(*n*+1)^|**y**)/*P*(θ^(*n*)^|**y**)>1, and it is accepted with probability *β* if *P*(θ^(*n*+1)^|**y**)/*P*(θ^(*n*)^|**y**) = *β* ≤1.

The algorithm described above was implemented using MATLAB R2020a, where the gLV models were solved using variable step solver ode23s. 120,000 MCMC samples were collected after a burn-in period of 10,000 samples. The Gelman-Rubin potential scale reduction factor (PSRF) was used to evaluate convergence of the posterior distribution estimates, where a PSRF closer to 1 indicates better convergence. The average PSRF is 1.31 and 80% of the parameters have a PSRF less than 1.5. The medians of the marginal distributions of all parameters correlated strongly with parameters estimated by the FMINCON method (Pearson r=0.99).

### Hill fits

The community sensitivity to *C. difficile* initial abundance was quantified by fitting the data to the Hill equation:

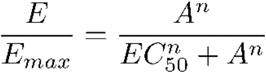

where *E* is 48 hour abundance of *C. difficile*, *E_max_* is the maximum 48 hour abundance of *C. difficile* across all initial fractions, *A* is the initial fraction of *C. difficile*, *EC_50_* is the initial fraction that produces 50% of *E_max_* value, and *n* is a measure of ultrasensitivity. The data was fit using custom python scripts implementing the curve_fit function of the scipy package optimization module.

### Normalized Euclidean Distances

The normalized Euclidean distance (D) between uninvaded resident community *R* and *C.* difficile-invaded community *V* is calculated using

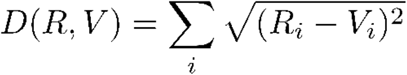

where *R* is the 48 hour timepoint of the uninvaded resident community and *V* is the 48 hour timepoint of the resident community invaded with *C. difficile*. *R_i_* is the relative abundance of species *i* in the uninvaded resident community, equal to reads of species *i* divided by the total community reads. *V_i_* is the normalized relative abundance of species *i* in the invaded community, equal to reads of species *i* divided by the resident community reads (total community reads minus *C. difficile* reads).

