## Supplementary Information for "Species richness determines *C. difficile* invasion outcome in synthetic human gut communities"

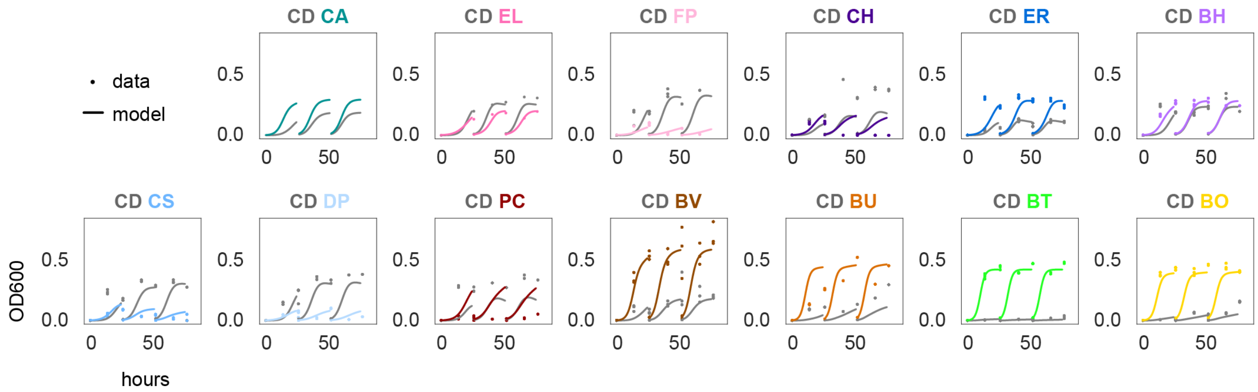


**Supplementary Figure S1: *C. difficile* introduced into pairwise communities with low initial density coexists with majority of resident gut species.** Absolute abundance (OD600) of species over time for three growth cycles. First growth cycle inoculated at a 1:9 ratio of *C. difficile* to resident species based on OD600 measurements. Datapoints indicate experimental biological replicates. Lines indicate simulations using the generalized Lotka-Volterra Full Model.

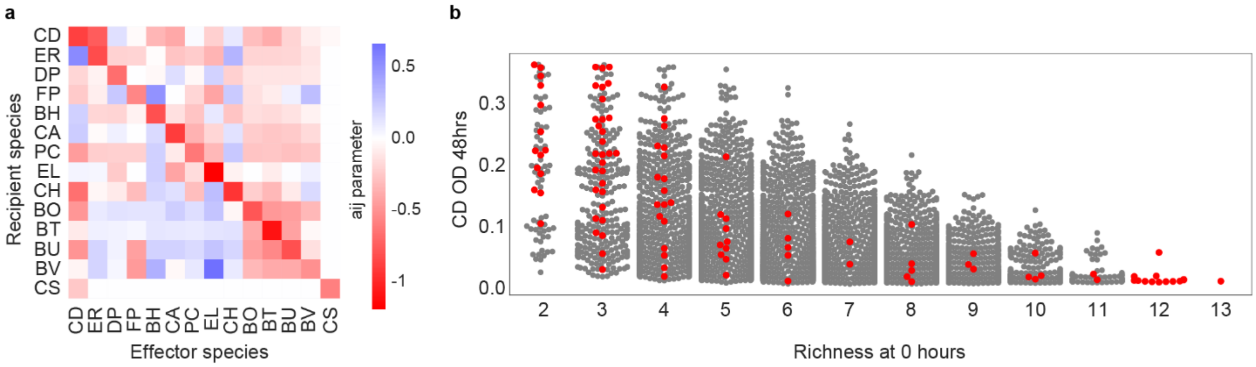


**Supplementary Figure S2: Generalized Lotka-Volterra model parameters for Preliminary Model and predictions of species abundance. (a)** Heatmap of inferred interspecies interaction parameters of Preliminary Model. **(b)** Swarmplot of simulated *C. difficile* absolute abundance (OD600) at 48 hours using the Preliminary model as a function of initial resident species richness for all 8,178 possible *C. difficile*-containing subcommunities composed of 2-13 species. Red points indicate communities chosen for experimental characterization.

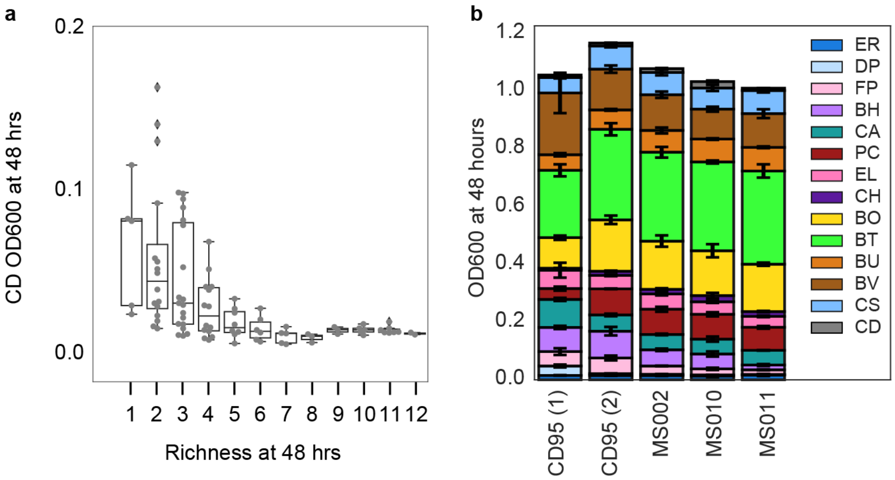


**Supplementary Figure S3: Analysis of 48-hour community composition of communities assembled with *C. difficile*. (a)** Swarmplot of *C. difficile* (CD) absolute abundance (OD600) at 48 hours in 94 sub-communities as a function of species richness at 48 hours. Datapoints indicate mean of two to three biological replicates. Line represents median, box edges represent first and third quartiles, and whiskers indicate the minimum and maximum. Outliers are denoted by diamonds. **(b)** Barplot of community composition of full community containing one of four different strains of *C. difficile* (**Table S1**). CD95 (1) condition had an initial total OD600 of 0.0066 and initial evenness of 1. CD95 (2) and MS conditions had an initial *C. difficile* OD600 of 0.00032 and initial non-*C. difficile* species OD600 of 0.00047. Bars indicate the mean and error bars indicate one standard deviation from the mean of three biological replicates.

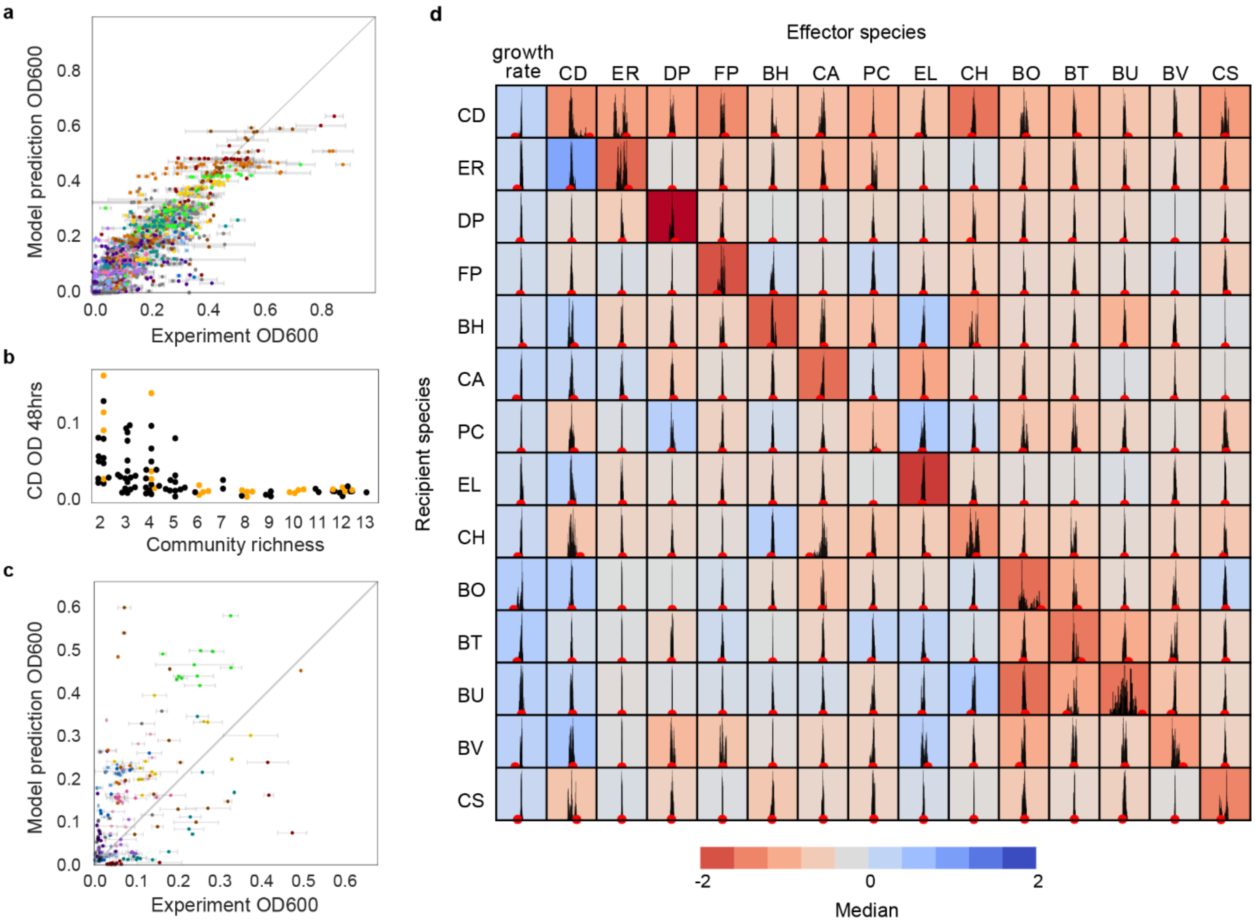


**Supplementary Figure S4**: **Analysis of parameter uncertainty and predictive capability of generalized Lotka-Volterra models.** **(a)** Scatterplot of mean experimental absolute abundance versus predicted species absolute abundance using the Full Model for the communities in the training data set (Pearson r=0.89, p<0.001). Error bars represent one standard deviation from the mean of two to three biological replicates. Gray line indicates y=x, or 100% prediction accuracy. **(b)** Swarmplot highlighting 24 communities chosen as the held-out set (data also shown in Fig. 2a). Orange datapoints represent held-out communities from training set. Black datapoints indicate communities from Fig 2a in training data set. **(c)** Scatterplot of average experimental absolute abundance (OD600) versus predicted species absolute abundance (OD600) using the Preliminary Model for the 24 held-out communities (Pearson r=0.84, p<0.001). Error bars represent one standard deviation from the mean of two to three of biological replicates. Gray line indicates y=x, or 100% prediction accuracy. **(d)** Histograms of parameter values determined using Metropolis-Hastings Monte Carlo (MCMC) analysis. Red dots indicate the parameter value in the Full Model (**Methods**). The black histograms indicate the MCMC distribution. The x-axis is scaled to median of MCMC distribution +/- 0.25. The color of each subplot denotes the median value of the MCMC distribution.

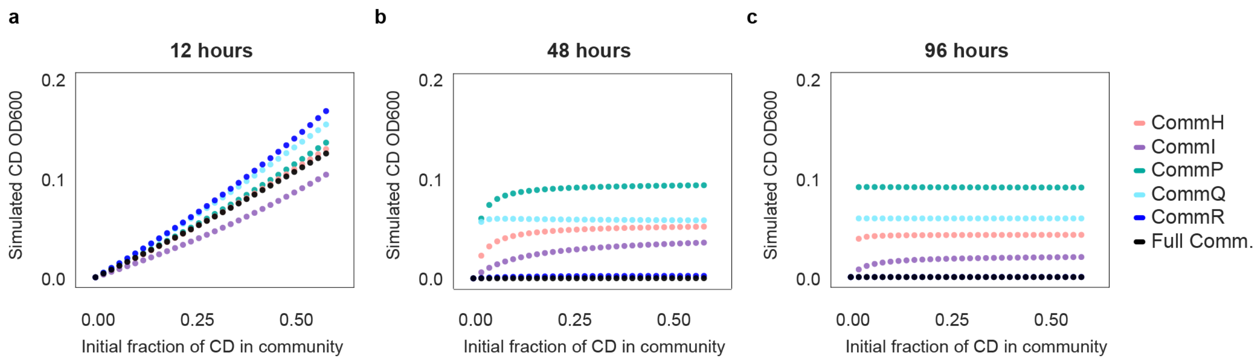


**Supplementary Figure S5: Predicted relationship between propagule pressure and *C. difficile* abundance at different time points.** Lineplots of predicted absolute abundance (OD600) of *C. difficile* using the Full Model in communities at **(a)** 12 hours, **(b)** 48 hours, and **(c)** 96 hours as a function of the initial fraction of *C. difficile*.

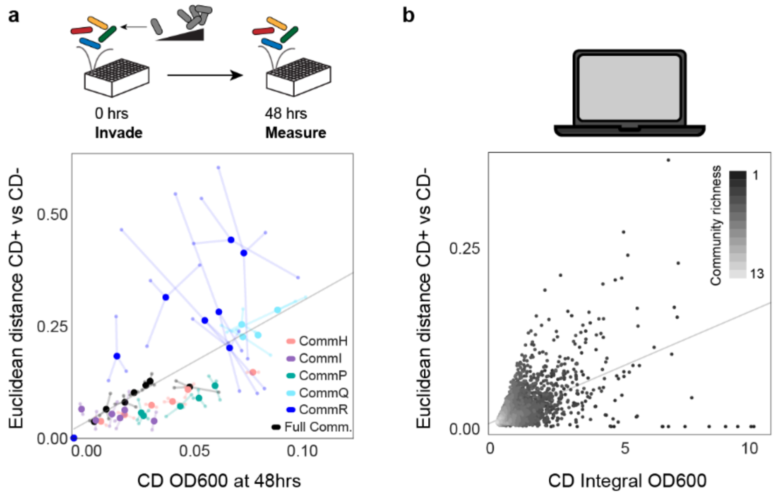


**Supplementary Figure S6: Change in resident community composition in the presence and absence of *C. difficile*. (a)** Scatterplot of normalized Euclidean distance (**Methods**) between communities initialized with various *C. difficile* abundances and the uninvaded communities as a function of *C. difficile* absolute abundance (OD600) in the community at 48 hours. Gray line indicates a linear regression (y=2.7x+0.02, Pearson r=0.61, $p<0.001$). Transparent data points indicate biological replicates and are connected to the corresponding mean values by transparent lines. **(b)** Scatterplot of normalized Euclidean distance between the 48 hour abundance of simulated communities invaded with *C. difficile* at six hours and uninvaded communities as a function of simulated *C. difficile* integral OD600 from 0 to 48 hours in the community (y=0.16x+0.004, Pearson r=0.58, $p<0.001$). Shading indicates species richness.

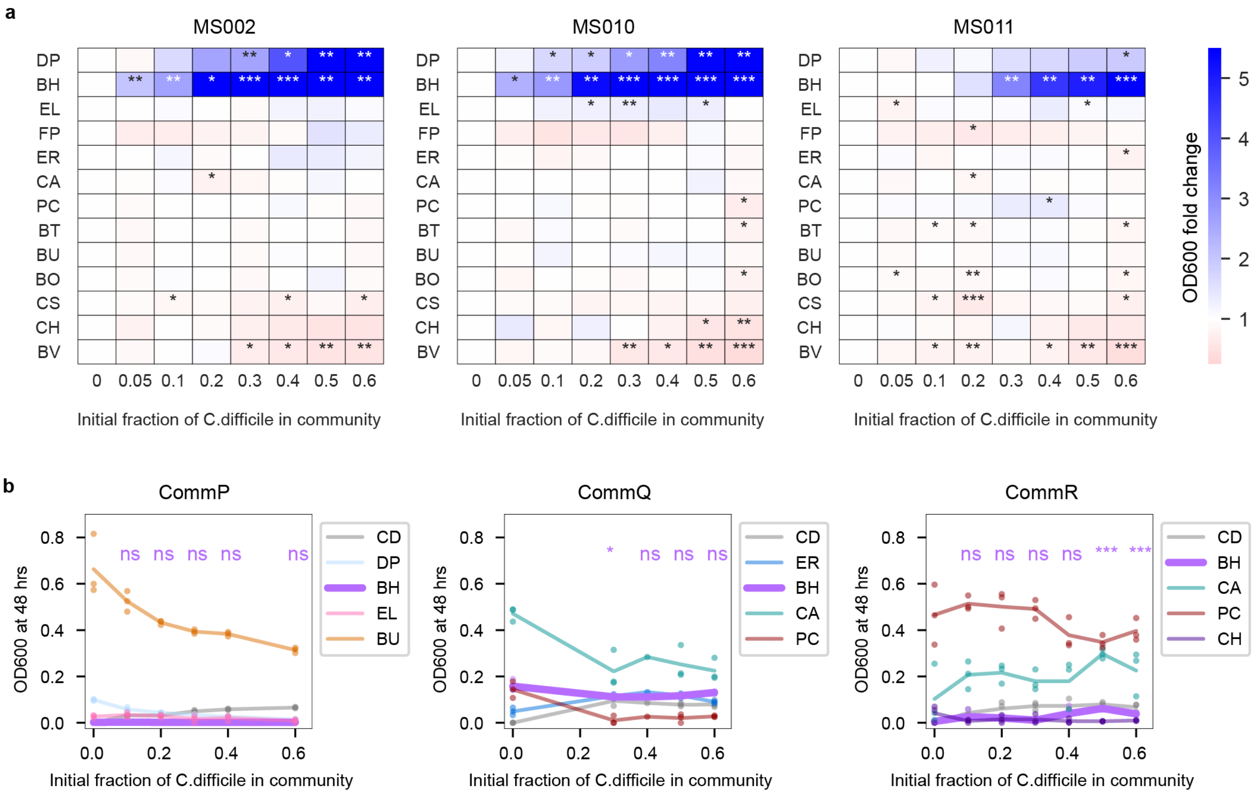


**Supplementary Figure S7: Impact of *C. difficile* on resident species abundances. (a)** Heatmap of the fold change of species absolute abundance (mean-value of three biological replicates) in full community with 5-60% initial *C. difficile* compared to the 0% initial *C. difficile* condition. Stars represent statistical significance: * p<0.05, ** p<0.01, *** p<0.001 according to an unpaired t-test. Left, middle, and right show communities with clinical *C. difficile* strains MS002, MS010, and MS011 respectively. **(b)** Lineplots of species absolute abundance (OD600) at 48 hours as a function of initial *C. difficile* fraction. Datapoints indicate biological replicates and lines indicate the mean. Stars indicate a statistically significant difference in the absolute abundance of *B. hydrogenotrophica* compared to the absolute abundance of *B. hydrogenotrophica* in 0% initial *C. difficile* condition: * p<0.05, ** p<0.01, *** p<0.001, ns=no significant difference according to an unpaired t-test. Left, middle, and right show communities CommP, CommQ, and CommR respectively.

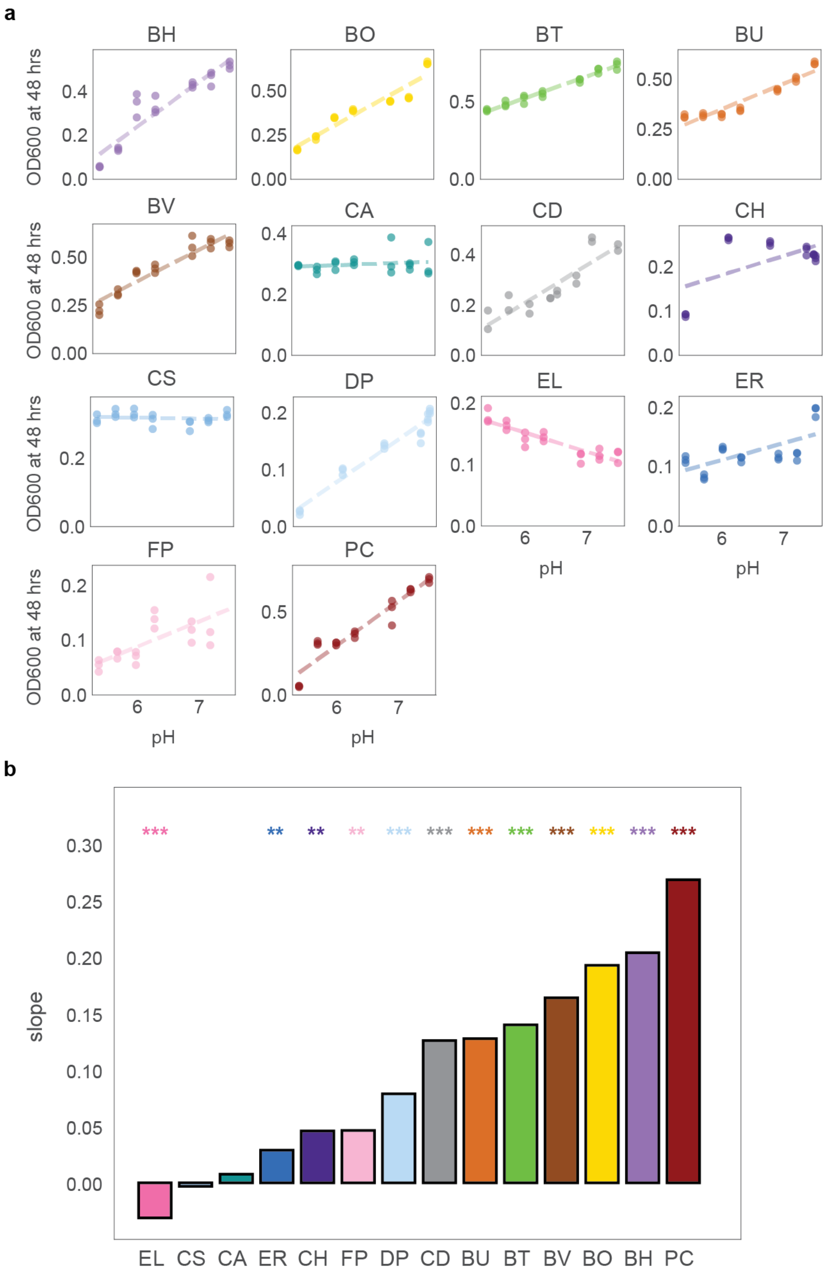


**Supplementary Figure S8: pH sensitivity of resident gut species in monoculture. (a)** Lineplots of monospecies absorbance at 600 nm (OD600) at 48 hours as a function of the initial environmental pH of adjusted fresh media. Datapoints indicate biological replicates and dashed lines indicate linear regression fits. **(b)** Barplot of slopes of linear regression fit to data in **a**. Stars denote statistical significance: * p<0.05, ** p<0.01, *** p<0.001 according to an unpaired t-test.

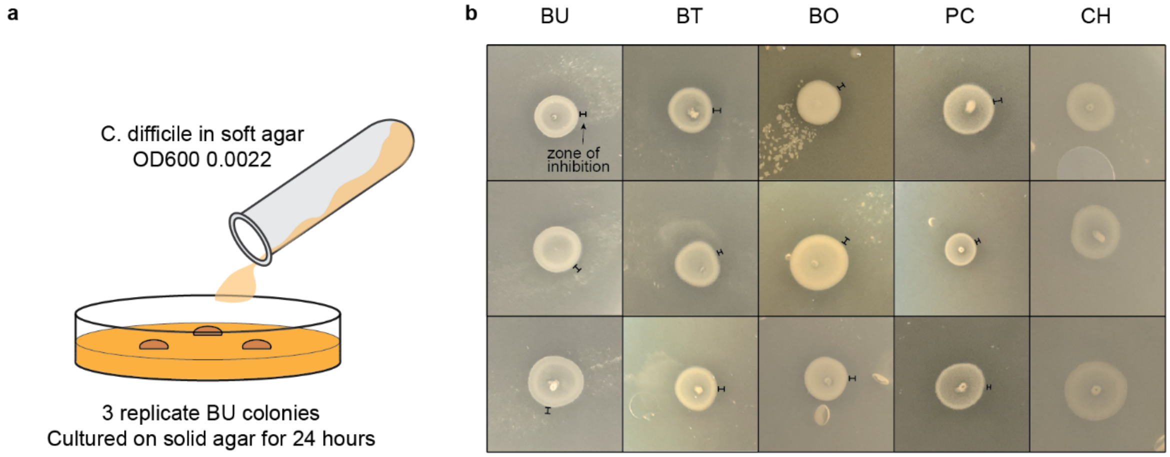


**Supplementary Figure S9: *C. difficile* growth inhibition in soft agar overlay assay. (a)** Illustration of soft agar overlay method. **(b)** Image of *C. difficile* growth in soft agar. Zones of inhibition at 24 hours were observed in 3 replicates for *B. uniformis, B. thetaiotaomicron, B. ovatus, P. copri*. No zones of inhibition were visible for *C. hiranonis*. Zones are marked with black bars.

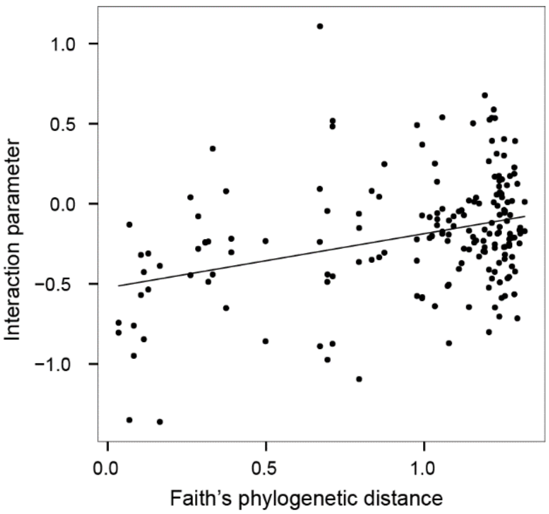


**Supplementary Figure S10**: **Interaction parameter increases with phylogenetic distance.** Scatterplot of interspecies interaction parameters ($\alpha_{ij}$s) from Full Model as a function of Faith's phylogenetic distance between species *i* and *j* (y=0.34x-0.52, Pearson r=0.34, p<0.001). Faith's phylogenetic distance was calculated from branch lengths of phylogenetic tree based on concatenated alignment of 37 marker genes.

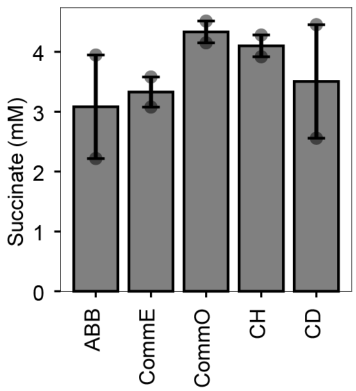


**Supplementary Figure S11: Succinate concentration in supernatants at 20 hours.** Barplot of succinate concentration determined by fluorometric enzymatic assay (**Methods**). Datapoints indicate technical replicates and bar indicates mean concentration. ABB denotes fresh media.
